# Structure-Activity Relationship Studies of Novel Gut-derived Lantibiotics Against Human Gut Commensals

**DOI:** 10.1101/2023.09.15.557961

**Authors:** Zhenrun J. Zhang, Chunyu Wu, Darian Dorantes, Téa Pappas, Anitha Sundararajan, Huaiying Lin, Eric G. Pamer, Wilfred A. van der Donk

## Abstract

Recent advances in sequencing techniques unveiled the vast potential of ribosomally synthesized and post-translationally modified peptides (RiPPs) encoded in microbiomes. Class I lantibiotics such as nisin A, widely used as a food preservative, have been investigated for their efficacy in killing pathogens. However, the impact of nisin and nisin-like class I lantibiotics on commensal bacteria residing in the human gut remains unclear. Here, we report six gut-derived class I lantibiotics that are close homologs of nisin, four of which are novel. We applied an improved lantibiotic expression platform to produce and purify these lantibiotics for antimicrobial assays. We determined their minimal inhibitory concentration (MIC) against both Gram-positive human pathogens and gut commensals, and profiled the lantibiotic resistance genes in these pathogens and commensals. SAR studies with variants revealed key regions and residues that impact their antimicrobial properties. Our characterization and SAR studies of nisin-like lantibiotics against both pathogens and human gut commensals could shed light on the future development of lantibiotic-based therapeutics and food preservatives.

## Introduction

Bacteria have evolved in intense competition with other microbes in complex environments. In order to make niche clearance or to establish colonization resistance in the community, many bacteria produce bacteriocins to kill microbial competitors.^1^ Bacteriocins can function as natural food preservatives through the inhibition of pathogenic bacteria, ultimately contributing to food safety.^2^ Ribosomally synthesized and post-translationally modified peptides (RiPPs) are a major class of bacteriocins that exhibit broad spectrum of antimicrobial activities against Gram-positive and Gram-negative bacteria including *S. aureus*, *E. faecalis* and *E. coli.*^3–5^ The biosynthesis of RiPPs starts with a gene-encoded precursor peptide, which comprises a C-terminal core peptide fused to an N-terminal leader peptide.^6^ The leader peptide is recognized by modification enzymes that are usually encoded in the same biosynthetic gene cluster (BGC) to receive post-translational modifications on the core peptide. Subsequently, the leader peptide is removed by protease cleavage to yield the mature peptide with desired bioactivity.^7^ Bioinformatic studies have found great potential for RiPP discovery from the microbiome.^8^ Recent advances in metagenomic sequencing and assembly make vast amounts of environmental microbial genomes available for investigation. Mining algorithms of BGCs with machine learning are also developed to accommodate rapidly increasing metagenomic datasets.^7, 9^ Lanthipeptides are a very large family of RiPPs that are characterized by thioether crosslinks called lanthionine and methyllanthionine.^10^ The bioinformatic tool RODEO (Rapid ORF Description & Evaluation Online)^11^ has been used to discover novel lanthipeptide families from more than one hundred thousand genomes.^12^

Lantibiotics, coined by combining “lanthipeptide” and “antibiotics”, are a class of RiPPs that have garnered significant attention due to their potent antimicrobial properties and wide application in the food industry. The thioether macrocycles in class I lantibiotics are installed in a two-step process.^10^ LanB dehydrates Ser and Thr residues in the LanA precursor peptides to generate dehydroalanine (Dha) and dehydrobutyrine (Dhb), respectively. The dehydration is followed by LanC catalyzing intramolecular Michael-type addition of Cys residues to Dha or Dhb, forming lanthionine or methyllanthionine. The catalytic activity of LanB depends on glutamyl-tRNA synthetase (GluRS) and tRNA^Glu^,^13^ and LanBs display sequence selectivity towards the tRNA^Glu^ acceptor stem.^14^ Previous studies have demonstrated that introducing GluRS and tRNA^Glu^ from the native lanthipeptide-producing organism could improve the production of fully modified peptides in *E. coli*.^14, 15^ Using this information, an improved production platform has been established for facile expression and production of class I lanthipeptides.^16^

Blauticin, a class I lantibiotic produced by the gut commensal bacteria *Blautia producta* SCSK (BP_SCSK_), exerts colonization resistance and clearance of Vancomycin-resistant *Enterococci* (VRE) *in vivo*.^17^ A previous study showed that the *in vivo* VRE colonization in colon was inhibited by BP_SCSK_ but not by *Lactococcus lactis*, the producing strain of the well-studied class I lantibiotic nisin. ^17^ In comparison to nisin, blauticin has reduced activity against intestinal commensal bacteria.^17^ Nisin binds lipid II and inhibits peptidoglycan biosynthesis through its N-terminal rings, and forms pores in the bacterial membrane that also involves the C terminus of the peptide.^18–20^ These pores are made up of eight nisin molecules and four lipid II molecules.^21^ Extensive efforts have been made to reveal the Structure-Activity Relationship (SAR) of nisin and its antimicrobial efficacy.^22-24^ However, the SAR of blauticin and its bioactivities against pathogens and human gut commensals remains to be elucidated.

The sensitivity of human gut commensals to nisin that is widely present in food^25^ has been surprisingly underexplored. Gram-positive human gut commensals, especially those within the Lachnospiraceae family of Bacillota (previously Firmicutes), are major producers of secondary metabolites such as short-chain fatty acids and secondary bile acids, which are crucial in contributing to the stability of the gut microbiome and host immune homeostasis.^26^ Orally ingested nisin A induced sizable but reversible changes in microbial composition and metabolic activities of gut microbiome in pigs^27^ and in mice.^28^ SAR studies of lantibiotics including blauticin and their potencies on human gut commensals will guide the future development of novel preservatives that could cause less collateral damage to human microbial communities. Genetic screening in pathogens and lantibiotic-producing organisms has identified multiple mechanisms of resistance against lantibiotics. These include cell wall modifications, cell membrane modifications, and efflux pumps, among others.^29^ However, whether these mechanisms exist and how they may impact lantibiotic resistance in human gut commensals remain to be studied.

Here, we applied RODEO to the public RefSeq database to uncover novel nisin-like class I lantibiotics encoded in gut microbial genomes. After going through filtering criteria, six gut-derived class I lantibiotics that are close homologs of nisin were discovered, four of which were novel. We applied the improved lantibiotic expression platform to produce and purify these lantibiotics for antimicrobial assays, and determined their minimal inhibitory concentration (MIC) against both Gram-positive human pathogens and gut commensals. Furthermore, we profiled the lantibiotic resistance genes in these pathogens and commensals. Detailed SAR studies with variants revealed key regions and residues in these lantibiotics that impact the antimicrobial properties. Our characterization and SAR studies of class I lantibiotics against both pathogens and human gut commensals could shed light on the future development of lantibiotic-based therapeutics and/or food preservatives.

## Results

### Mining of class I lantibiotics from the gut

To find novel class I lantibiotic sequences from the gut, we adopted a workflow shown in Fig. 1A. We applied RODEO,^11^ an algorithm previously described to identify lanthipeptide sequences in large numbers of genomes,^12^ to the RefSeq database. RODEO uses a hidden Markov model to mine for RiPP BGCs and predict precursor peptides by combination of heuristic scoring and machine learning.^11^ We restricted our search to bacterial genomes whose representatives were reported to be found in mammalian guts. From this approach, we identified 41 unique lanthipeptide candidates (Supporting Information). To find close homologs of nisin and related lantibiotics, we filtered the results based on the following criteria. First, the peptide length had to be smaller or equal to 72. Second, the cysteine positions in the peptide needed to align with the ring-forming Cys residues in nisin A and blauticin based on a multiple sequence alignment (Fig. S1). We further consolidated sequences that differed only in leader sequences but were the same in the core peptide. Six unique class I lantibiotic sequences, including two known examples, blauticin from *Blautia producta* and nisin O from *Blautia obeum,*^30^ were identified (Fig. 1B, Supporting Information). In fact, blauticin was also found in *Blautia coccoides* and nisin O was also found within a *Faecalicatena contorta* genome. The sequences of four novel nisin-like class I lantibiotic candidates were compared to the prototypical lantibiotics nisin A and blauticin, as shown in Fig. 1B. Lan-Df from *Dorea formicigenerans* was identified to be a close homolog of nisin O, with one amino acid different from nisin O at ring A. Lan-CE02 from *Clostridium* sp. E02 further deviates from the prototype lantibiotics, with unique residues spanning across the peptide. We discovered two variants, Lan-P49.1 and Lan-P49.2, from the same genome of *Pseudobutyrivibrio* sp. 49. Lan-P49.1 and Lan-P49.2 both share the same sequences as blauticin in ring A and B at the N termini of the peptides, but the C-termini of the peptides including the hinge region between rings C and D are different from nisin A and blauticin. This hinge region has been shown to be important for pore formation and antimicrobial activity.^31–34^

**Fig. 1.**
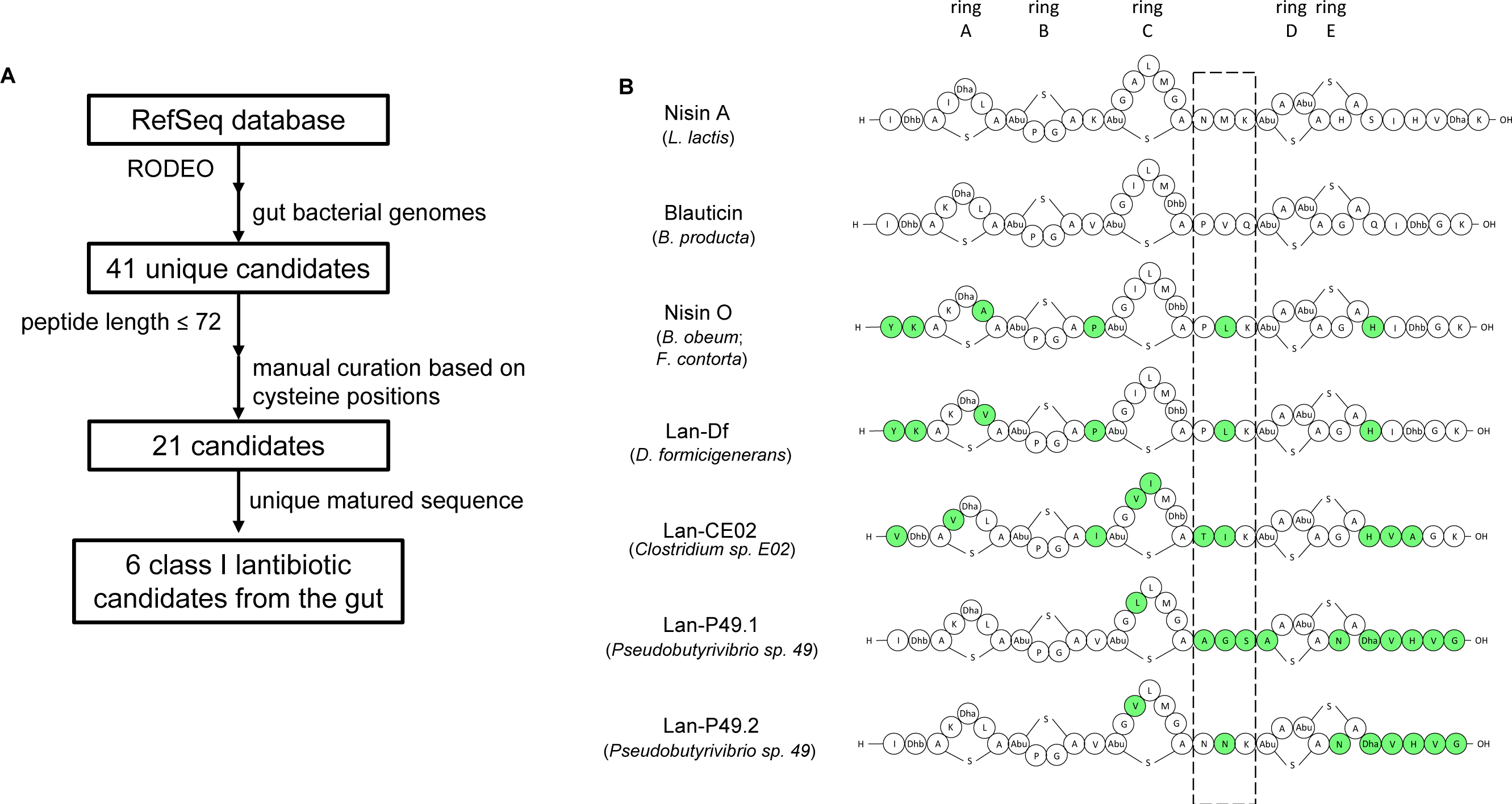
Bioinformatic discovery of nisin-like class I lantibiotics encoded in gut bacteria. **(A)** Workflow chart of lantibiotic discovery from RefSeq database using RODEO and filtering criteria. **(B)** Illustration of discovered putative class I lantibiotics with the ring patterns predicted based on that of nisin A. Green shading indicates residues different from both nisin A and blauticin. Dashed box indicates hinge region. Ring designations in the structure are denoted on the top.

### Heterologous expression of blauticin in *E. coli*

The genome of *Blautia producta* SCSK has been sequenced, assembled and annotated.^35^ To heterologously express blauticin in *E. coli*, a three-plasmid-based expression platform that was reported previously was applied (Fig. S2A).^16^ In the current version, this platform includes a pETDuet plasmid that encodes a His-tagged blauticin leader and core sequence. The blauticin leader sequence is recognized by blauticin-specific LanB and LanC (*bpcB* and *bpcC* gene, respectively) encoded in the pCDFDuet plasmid. A third pEVOL plasmid encodes two copies of glutamyl-tRNA synthetase (GluRS) from *B. producta* (one under an inducible promoter araBAD, the other under a constitutive promoter glnS) and a copy of tRNA^Glu^ from *B. producta* under a constitutive promoter proK. The availability of cognate glutamyl-tRNA was envisioned to ensure optimal catalytic activity of BpcB as the sequence of tRNA^Glu^ strongly affects LanB dehydration activity.^14^ Coding sequences of *bpcB* and *bpcC* genes were codon-optimized for expression in *E. coli* (Supporting Information). Heterologously expressed lantibiotic precursors were purified from cell lysate by metal affinity chromatography, followed by trypsin digestion for leader peptide removal and RP-HPLC purification (Fig. S2B).

Among the five BP_SCSK_ lantibiotic precursors that are encoded in the blauticin BGC, the BP_SCSK_ lantibiotic precursors BpcA_1_-BpcA_4_ are four identical copies of the blauticin precursor peptide and BpcA_5_ is a fifth peptide the function of which remains to be identified (Fig. 2A).^17^ An initial attempt of co-expressing His_6_-tagged BpcA_1_ with BpcB and BpcC in *E. coli* in the absence of the pEVOL plasmid resulted in a mixture of zero to nine dehydrations with the nine-fold dehydrated peptide as one of the least abundant products after leader peptide removal (Fig. 2D). The poor dehydratase activity is likely the result of sequence differences of the major recognition elements for the lantibiotic dehydratases that are located on the acceptor stem of tRNA^Glu^ when comparing *E. coli* and *B. producta* sequences (Fig. 2B). After incorporating the pEVOL vector that encodes *B. producta* SCSK GluRS and tRNA^Glu^_CUC_ in the expression system, up to nine dehydrations were observed by matrix-assisted laser desorption ionization time-of-flight mass spectrometry (MALDI-TOF MS) after trypsin removal of the leader peptide, and the mass distribution of the product looked similar as that of wild-type blauticin that was isolated from the producing organism (Fig. 2C and E). Production of lantibiotics as a mixture of peptides with different dehydration states in the native producing organism is not uncommon and is for example also observed for the commercial food preservative nisin ^36^ or a recently reported lanthipeptide from the human oral microbiome that has pro-immune activity.^37^ We further purified the blauticin mixture with RP-HPLC and collected its fully dehydrated form (nine dehydrations) and compared its antimicrobial activity to the dehydration mixture (Fig. S4A). We observed similar minimal inhibitory concentration (MIC) of fully dehydrated blauticin, mixed dehydrated blauticin, and native blauticin against Vancomycin-resistant *Enterococcus faecium* ATCC700221 (VRE) (Fig. S4B).

**Fig. 2.**
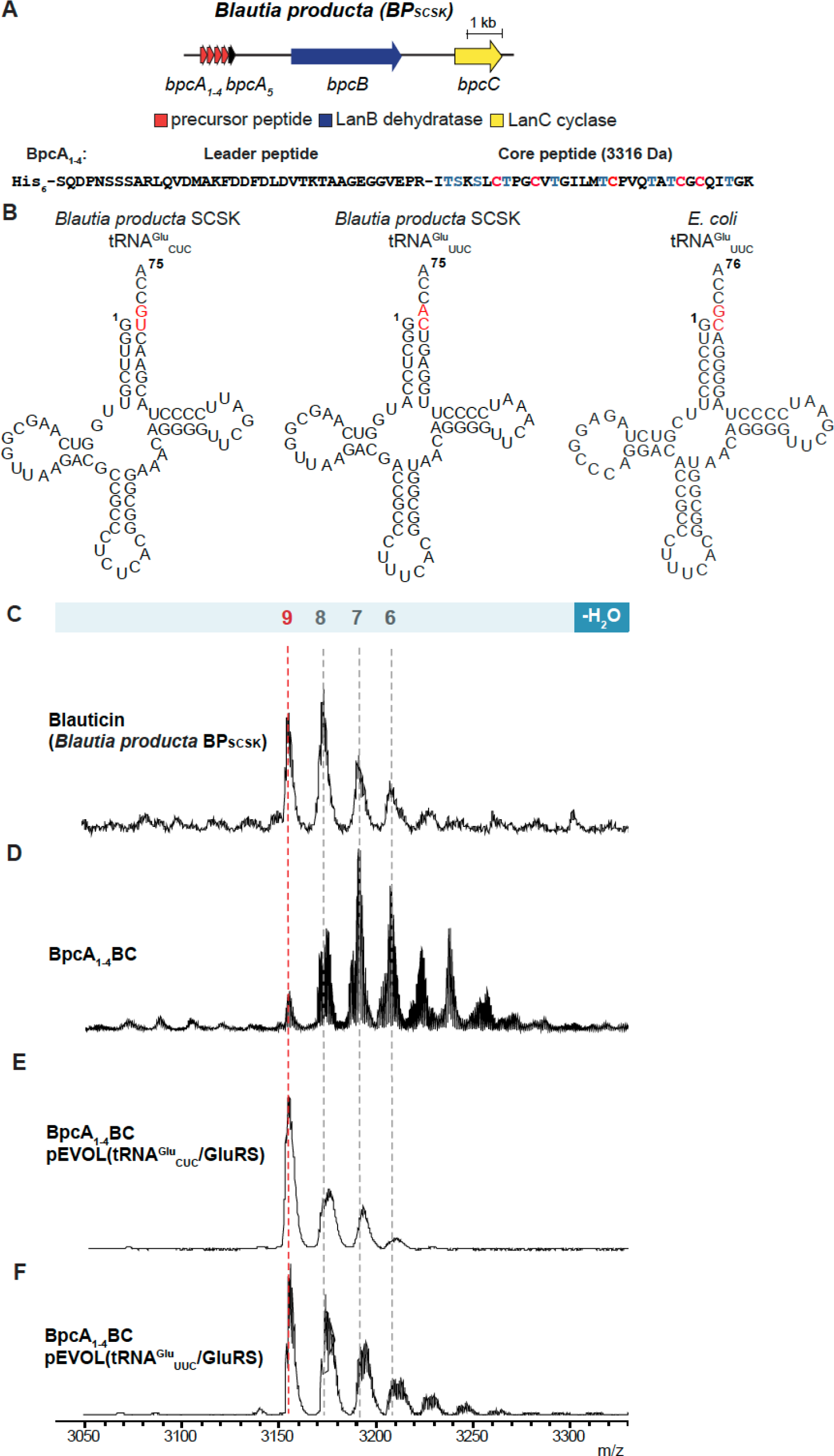
Heterologous production of blauticin using the pEVOL vector in *E. coli* and leader peptide removal with trypsin. **(A)** Blauticin BGC from *B. producta* SCSK. **(B)** Predicted cloverleaf structures of *B. producta* SCSK and *E. coli* tRNA^Glu^ made with the tRNAscan-SE algorithm.^38, 39^ The proposed major recognition elements of the tRNA^Glu^ acceptor stem used by lantibiotic dehydratases are highlighted in red. **(C-F)** MALDI-TOF MS analysis of blauticin isolated from BP_SCSK_ (C), His_6_-BpcA_1_ isolated from the co-expression with BpcBC without (D) and with the pEVOL platform using tRNA^Glu^_CUC_ (E) or tRNA^Glu^_UUC_ (F) in *E. coli* after leader peptide removal by trypsin. 6, 6-fold dehydrated BpcA_1_ ([M + H]^+^ m/z 3,210, calc. m/z 3,208); 7, 7-fold dehydrated BpcA_1_ ([M + H]^+^ m/z 3,192, calc. m/z 3,190); 8, 8-fold dehydrated BpcA_1_ ([M + H]^+^ m/z 3,175, calc. m/z 3,172); 9, 9-fold dehydrated BpcA_1_ ([M + H]^+^ m/z 3,156, calc. m/z 3,154).

A previous report studied the tRNA specificity of the lantibiotic dehydratase MibB involved in the biosynthesis of NAI-107 from the Actinobacterium *Microbispora* sp. 107891.^14^ MibB was shown to not only discriminate between tRNA^Glu^ from different organisms, but also discriminate between tRNA^Glu^ isoacceptors encoded in the genome of *Microbispora* sp. 107891. A tRNA scanning analysis of the *B. producta* SCSK genome revealed the presence of two tRNA^Glu^ genes encoding tRNA^Glu^_CUC_ and tRNA^Glu^_UUC_ isoacceptors (Fig. 2B).^38, 39^ The presence of the two different tRNA^Glu^ types prompted investigation whether the dehydratase BpcB displayed isoacceptor preference. We co-expressed each of the two isoacceptors in the pEVOL vector with BpcA_1_, BpcB, and BpcC in *E. coli*. MALDI-TOF MS analysis of dehydration assays revealed no significant differences of BpcA_1_ dehydration (Fig. 2E and F), indicating that both tRNA^Glu^ isoacceptors can be accepted by BpcB.

### Isolation and bioactivities of novel class I lantibiotics

After successful demonstration of expression of blauticin in *E. coli*, we applied a similar approach to express and isolate the novel nisin-like class I lantibiotics discovered in the genomes of mammalian gut microbiota. Rather than using the precursor peptide and biosynthetic enzymes encoded in the producing organisms, we used the His-tagged blauticin leader sequence and fused to its C-terminus the desired lanthipeptide core sequence. The pETDuet plasmid encoding the chimeric substrate was co-expressed with plasmids encoding BpcB, BpcC, and *B. producta* GluRS and tRNA^Glu^. Such use of one set of producing enzymes for production of structurally closely related analogs of a certain lantibiotic has been successfully used previously.^40^ The modified lantibiotic precursor peptides were purified and digested with trypsin for leader peptide removal. Fully dehydrated Lan-CE02, Lan-Df, Lan-P49.1 and Lan-P49.2 were observed by MALDI-TOF MS (Figure S3 and Table S1).

Next, we set out to test the antimicrobial activities of these purified lantibiotics against a panel of Gram-positive bacteria that are representatives of some of the most prevalent human pathogens: VRE, Methicillin-resistant *Staphylococcus aureus* USA300 (MRSA), Methicillin-resistant *Staphylococcus epidermidis* SK135 (MRSE), *Listeria monocytogenes* 10403S, and *Clostridioides difficile* VPI10463. We examined the growth of bacteria in a 96-well plate under a concentration gradient of lantibiotics under anaerobic conditions (Fig. S2B). In general, nisin A and Lan-Df had the strongest inhibitory activities against pathogens based on their MIC, followed by nisin O. Blauticin, Lan-CE02, and Lan-P49.2 had intermediate activities, while Lan-P49.1 had the weakest activity (Fig. 3A-E). Comparing across the pathogen panel, VRE and *C. difficile* were more sensitive to lantibiotic treatment, while *L. monocytogenes* was the most resistant amongst the panel tested.

**Fig. 3.**
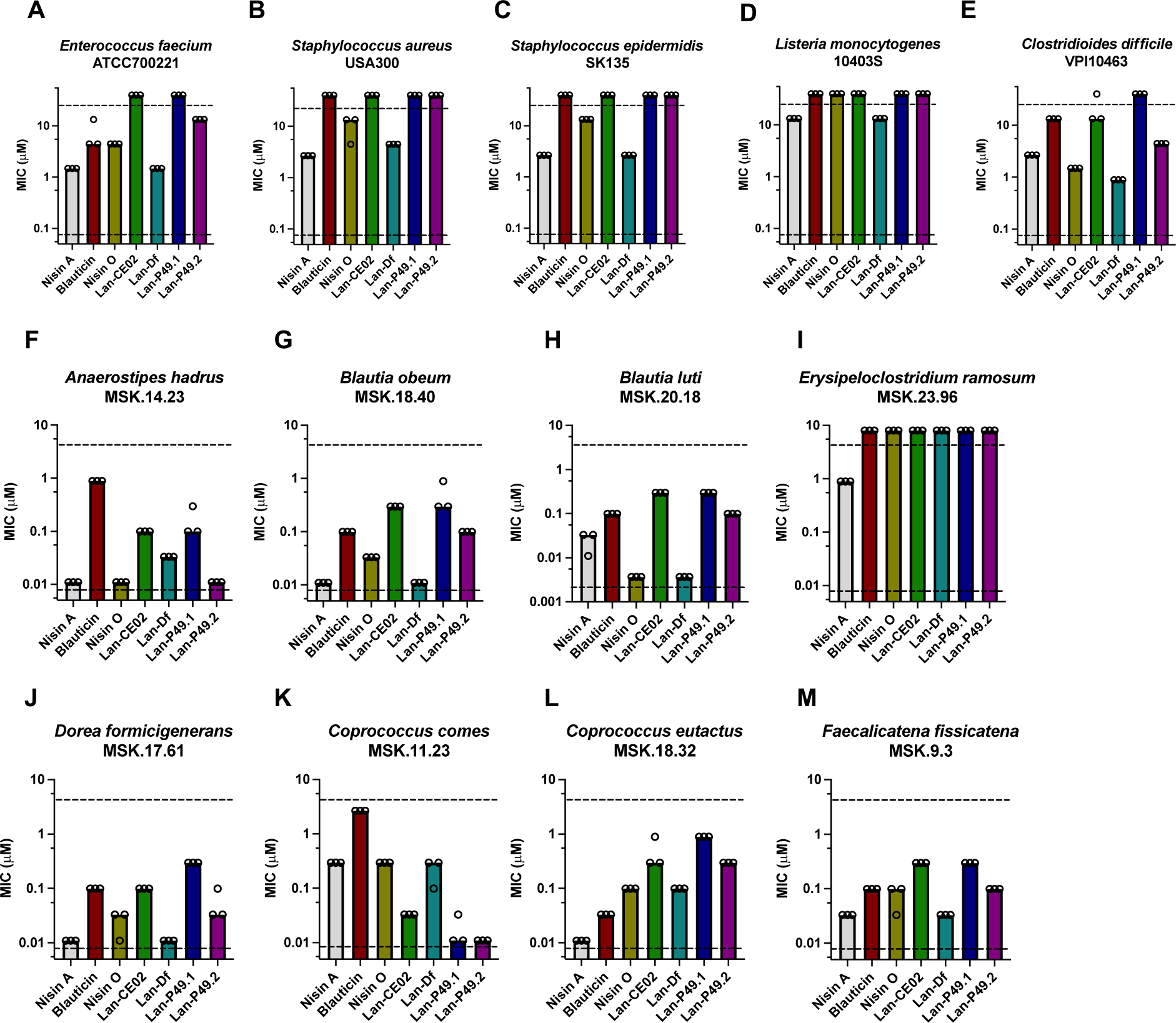
Minimal Inhibitory Concentration (MIC) of the panel of lantibiotics against human pathogens (A-E) and human gut commensals (F-M). Each dot indicates one measurement of MIC value. Each bar indicates median MIC value. Upper and lower dashed lines indicate upper and lower limit of concentration tested, respectively.

Antibiotic treatment may result in collateral damage to commensal bacteria in the human microbiome, leading to dysbiosis and susceptibility to various diseases.^41^ How lantibiotics impact Gram-positive commensals from the human gut has remained unclear. Therefore, we performed antimicrobial activity tests of these lantibiotics against select gram-positive human gut commensals, most of which belong to the Lachnospiraceae family, one of the most abundant bacterial families in the human gut.^26^ In general, human gut commensals are more susceptible to the set of investigated lantibiotics compared to pathogens (Fig. 3). The activities of the lantibiotics tested generally followed the trend of those against Gram-positive pathogens; while nisin A and Lan-Df were generally more effective in growth inhibition, Lan-P49.1, Lan-P49.2 and Lan-CE02 had weaker antimicrobial activities (Fig. 3F-M). Amongst all, *Erysipeloclostridium ramosum* MSK.23.96 displayed the highest resistance to lantibiotics compared to other commensals tested. Interestingly, *Coprococcus comes* MSK.11.23 had an opposite sensitivity profile; it was relatively resistant to nisin A, nisin O and Lan-Df, while it was sensitive to Lan-P49.1 and LanP49.2.

### Lantibiotic resistance genes in pathogens and commensals

The differences in lantibiotic susceptibility profiles of pathogens and human gut commensals lead to the further investigation of potential lantibiotic resistance mechanisms encoded in the genomes of gut bacteria. Many mechanisms of lantibiotic resistance in bacteria have been reported, mostly revealed through genetic screening of pathogens. One major resistance mechanism is bacterial cell wall modifications. The *dltABCD* operon is responsible for the D-alanylation of lipoteichoic acids (LTA) and wall teichoic acids (WTA), resulting in an increase of positive charges in the cell wall and the repulsion of cationic lantibiotics.^42, 43^ Lipid II, the target of nisin-like lantibiotics, is synthesized by MurG, flipped to the outer membrane by MurJ,^44, 45^ and used by penicillin-binding proteins (PBPs) as substrates for peptidoglycan biosynthesis.^46^ Therefore, these genes have been implicated in lantibiotic resistance.^47^ Another major resistance mechanism is efflux pumps. The LanFEG three-component transporter and homologs can expel lantibiotics,^48, 49^ while the BceAB two-component transporter, originally reported as a bacitracin efflux pump, also provides cross-protection against lantibiotics.^50^ Other resistance mechanisms include the tellurite resistance gene *telA*,^51^ cell membrane modifications by *mprF*,^52^ the nisin resistance protein NSR,^53, 54^ and lantibiotic self-resistance proteins LanI.^55^ Of these LanFEG and LanI are often found in lantibiotic BGCs.

We performed a Hidden Markov Model and BLAST search for the aforementioned genes among the genomes of pathogens and human gut commensals tested. For resistance mechanisms that require multiple genes (*dltABCD, bceAB* and *lanFEG*), only the complete presence as an operon was counted. The numbers of corresponding genes in each genome are listed in Tables 1 and 2. Bacteria are listed in ascending order from left to right according to the median MIC value across the nisin-like lantibiotic panel. We found that for most organisms, the numbers of potential resistance-inducing genes present in a specific genome did not correlate with stronger lantibiotic resistance (lower susceptibility). *becAB* and *mprF* may correlate with resistance in some cases, as the most sensitive *C. difficile* does not possess these genes. Among human gut commensals, the most resistant *E. ramosum* has more *bceAB* genes than others, and ties most for PBPs. However, one of the most sensitive strains, *A. hadrus* MSK.14.23, also contains *bceAB* genes, and ties most for PBPs. In conclusion, the numbers of putative resistance genes in these pathogens and commensals cannot fully explain their relative susceptibility against the lantibiotics tested.

**Table 1.** Number of genes in pathogens related to resistance to class I lantibiotics.

| Bacterial strain | *C. difficile*<br>VPI10463 | *E. faecium*<br>ATCC700221 | *S. aureus*<br>USA300 | *S. epidermidis*<br>SK135 | *L. monocytogenes*<br>10403s | Mechanism of resistance |
| --- | --- | --- | --- | --- | --- | --- |
| Nisin A MIC (μM) | 2.67 | 1.48 | 2.67 | 2.67 | 2.67 |  |
| Median MIC (μM) | 4.44 | 13.3 | >13.3 | >13.3 | >13.3 |  |
| Cell wall biosynthesis |  |  |  |  |  |  |
| *murG* | 1 | 2 | 1 | 1 | 1 | Biosynthesis of Lipid II |
| *murJ* | 4 | 3 | 1 | 2 | 3 | Lipid II flippase |
| Penicillin-binding proteins | 5 | 8 | 6 | 5 | 6 | Peptidoglycan synthesis from Lipid II |
| Complete *dltABCD* | 1 | 1 | 1 | 1 | 1 | Changing cell wall charge |
| Efflux pump |  |  |  |  |  |  |
| Complete *lanFEG* | 1 | 0 | 1 | 0 | 0 | Lantibiotic efflux |
| Complete *bceAB* | 0 | 1 | 1 | 1 | 1 | Bacitracin efflux |
| Others |  |  |  |  |  |  |
| *telA* | 1 | 1 | 1 | 1 | 1 | Unknown |
| *mprF* | 0 | 2 | 1 | 1 | 1 | Changing cell membrane charge |
| *nsr* | 0 | 0 | 0 | 0 | 0 | Lantibiotic protease |
| *lanI* | 0 | 0 | 0 | 0 | 0 | Lantibiotic sequestration |

**Table 2.** Number of genes in human gut commensals related to resistance of nisin-like lantibiotics.

| Bacterial strain | *A. hadrus*<br>MSK.14.23 | *D. formicigenerans*<br>MSK.17.61 | *B. obeum*<br>MSK.18.40 | *C. eutactus*<br>MSK.18.32 | *B. luti*<br>MSK.20.18 | *F. fissicatena*<br>MSK.9.3 | *C. comes*<br>MSK.11.23 | *E. ramosum*<br>MSK.23.96 | Mechanism of resistance |
| --- | --- | --- | --- | --- | --- | --- | --- | --- | --- |
| Nisin A MIC (μM) | 0.0110 | 0.0110 | 0.0110 | 0.0110 | 0.0329 | 0.0329 | 0.296 | 0.889 | |
| Median MIC (μM) | 0.0329 | 0.0329 | 0.0988 | 0.0988 | 0.0988 | 0.0988 | 0.296 | >2.67 | |
| Cell wall biosynthesis |  |  |  |  |  |  |  |  |  |
| *murG* | 1 | 1 | 1 | 1 | 1 | 1 | 1 | 1 | Biosynthesis of Lipid II |
| *murJ* | 2 | 2 | 4 | 4 | 4 | 2 | 4 | 2 | Lipid II flippase |
| Penicillin-binding proteins | 9 | 7 | 7 | 6 | 6 | 6 | 8 | 9 | Peptidoglycan synthesis from Lipid II |
| Complete *dltABCD* | 0 | 0 | 0 | 0 | 0 | 0 | 0 | 0 | Changing cell wall charge |
| Efflux pump |  |  |  |  |  |  |  |  |  |
| Complete *lanFEG* | 0 | 0 | 0 | 0 | 0 | 0 | 0 | 0 | Lantibiotic efflux |
| Complete *bceAB* | 1 | 0 | 0 | 1 | 0 | 0 | 1 | 2 | Bacitracin efflux |
| Others |  |  |  |  |  |  |  |  |  |
| *telA* | 0 | 0 | 1 | 0 | 1 | 1 | 1 | 0 | Unknown |
| *mprF* | 0 | 0 | 0 | 0 | 0 | 0 | 0 | 0 | Changing cell membrane charge |
| *nsr* | 0 | 0 | 0 | 0 | 0 | 0 | 0 | 0 | Lantibiotic protease |
| *lanI* | 0 | 0 | 0 | 0 | 0 | 0 | 0 | 0 | Lantibiotic sequestration |

### Structure-Activity Relationship Studies

Given the high similarity of sequences among the nisin-like lantibiotics encoded in the gut microbiome, we next set out to study Structure-Activity Relationships (SAR) between specific residues and the antimicrobial activity. Blauticin and nisin O are two lantibiotics derived from *Blautia* species, but their activities vary against both pathogens and commensals, with nisin O being more effective in most cases. Four major regions differ between them: residues 1 and 2 at their N termini, residue 12 between rings B and C, residues 20-22 at the hinge region, and residue 29 at the C termini (Fig. 4A). To assess the impact of these individual regions on the antimicrobial activity, we synthesized four blauticin analogs by replacing the blauticin residues with the corresponding residues in nisin O (SAR1 to SAR4) using the expression platform described above (Fig. 4A, Fig. S5 and Table S1).

**Fig. 4.**
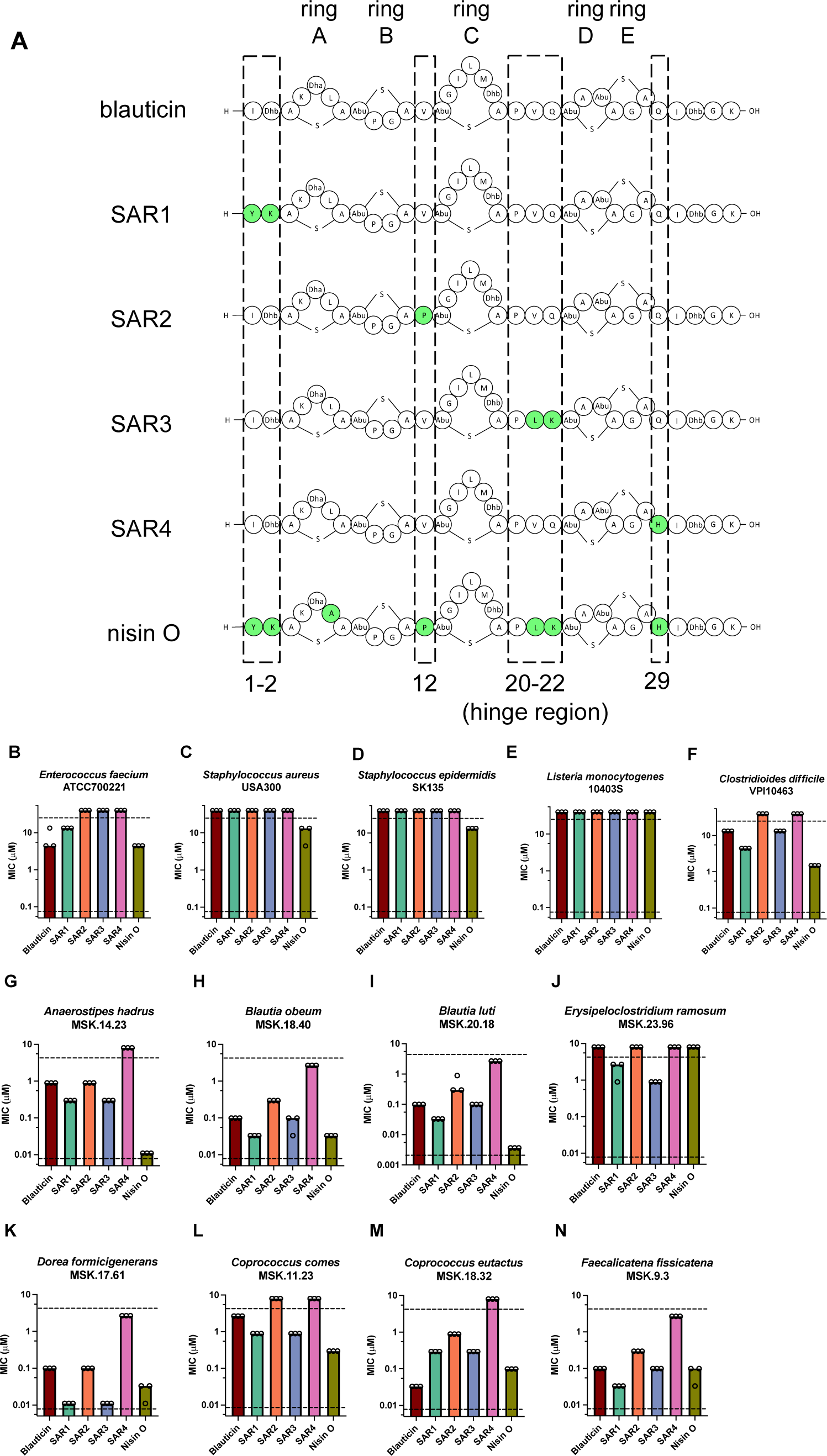
Structure-Activity Relationship (SAR) study of blauticin and comparison with Nisin O. **(A)** Illustration of blauticin, SAR variants and nisin O. Dashed boxes indicate regions from nisin O swapped into the blauticin backbone individually that created each SAR variant. Green shading indicates residues different from blauticin. Five rings in the structure are denoted on the top. Residue numbers and hinge region are denoted at the bottom. **(B-N)** Minimal Inhibitory Concentration (MIC) of the lantibiotic variants against human pathogens **(B-F)** and human gut commensals **(G-N)**. Each dot indicates one measurement of MIC value. Each bar indicates median MIC value. Upper and lower dashed lines indicate upper and lower limit of concentration tested, respectively. Observed and expected masses for each compound are listed in Table S1.

The bioactivities of the four variants were tested against the aforementioned panel of pathogens and gut commensals. In the pathogen panel, SAR1 generally had intermediate antimicrobial activity between blauticin and nisin O (Fig. 4B-F), while SAR2 to SAR4 displayed comparable or reduced activities compared to the two parent lantibiotics. On the other hand, for the gut commensal panel, SAR1 and SAR3 had intermediate inhibitory activities between blauticin and nisin O for some commensals (Fig. 4G-I, Fig. 4L), and had even better antimicrobial activities when targeting *E. ramosum* (Fig. 4J) or *D. formicigenerans* (Fig. 4K). On the contrary, SAR2 and SAR4 exhibited diminished activities compared to both parent molecules (Fig. 4G-N). Interestingly, for *C. eutactus*, none of the new variants displayed better inhibition than either parent lantibiotic, and it was also the only case when blauticin had a lower MIC than nisin O. These data suggest that the first two residues and the hinge region of nisin-like lantibiotics have more direct impact on the antimicrobial activity of nisin-like lantibiotics, while single-residue mutations V12P and Q29H had a deleterious impact on antimicrobial activity depending on the context of the backbone.

We further performed SAR studies on two highly similar novel lantibiotics, Lan-P49.1 and Lan-P49.2. They originate from the same *Pseudobutyrivibrio* genome, and their amino acid sequences have the largest difference in the hinge region. To test the impact of the hinge region, we swapped the hinge region of Lan-P49.1 into the Lan-P49.2 backbone, creating SAR5 (Fig. 5A, Fig. S5 and Table S1). SAR5 had decreased antimicrobial activity against the pathogen panel compared to Lan-P49.2 (Fig. 5B-F). When tested against the gut commensal panel, SAR5 displayed intermediate activity compared to Lan-P49.1 and Lan-P49.2 in most cases (Fig. 5H,I,K,M,N), reaching the same MIC as that of Lan-P49.1 in some bacteria (Fig. 5M) and the same MIC as that of Lan-P49.2 in other commensals (Fig. 5H,I,N). Interestingly, for *A. hadrus* and *C. comes*, SAR5 had decreased antimicrobial activity compared to both parent lantibiotics, whereas for *E. ramosum*, SAR5 had increased antimicrobial activity. These data demonstrate the importance of the hinge region in determining the antimicrobial activity of nisin-like lantibiotics, and are consistent with an emerging realization that lantibiotics can act differently based on the target organism investigated.^56^

**Fig. 5.**
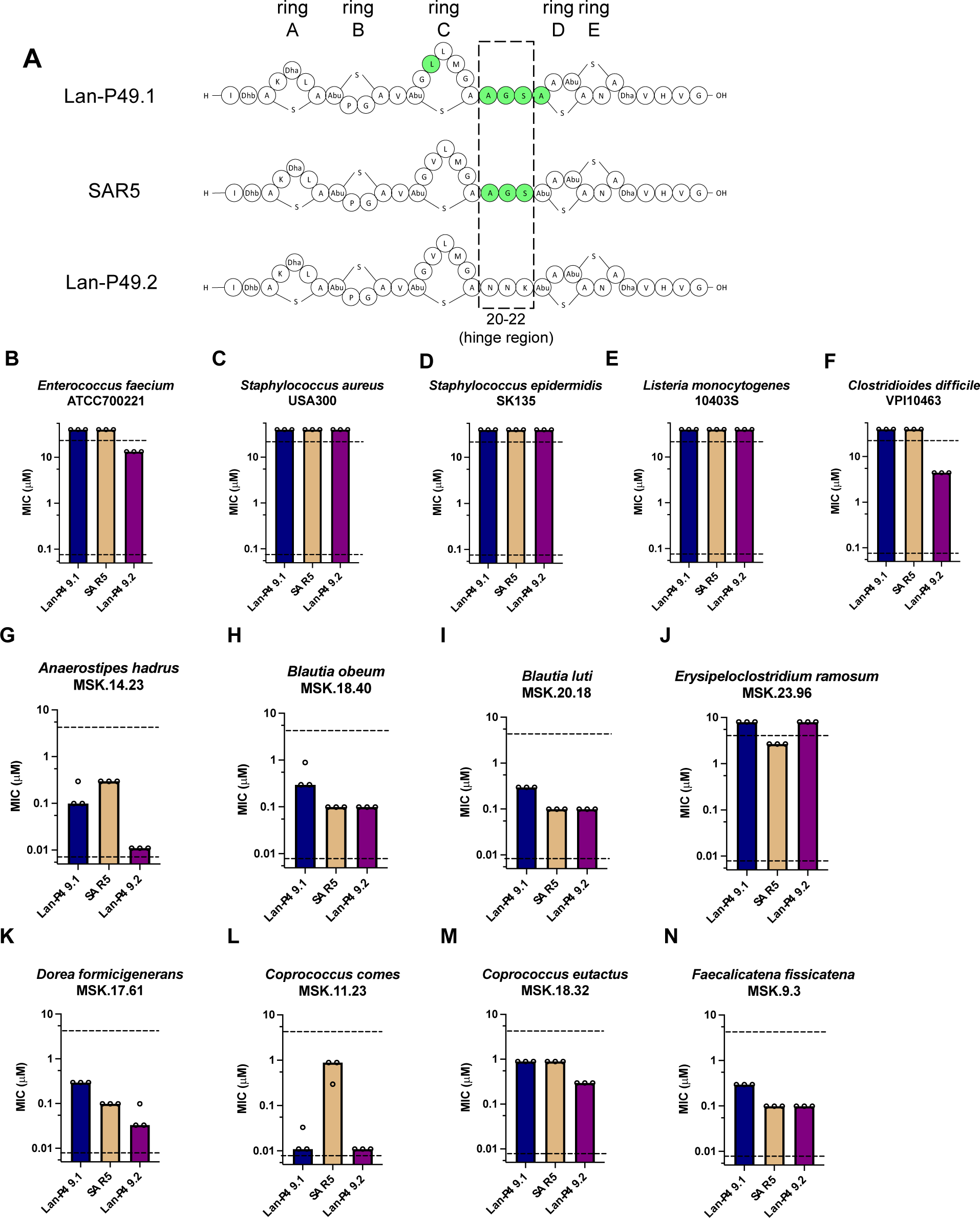
Structure-Activity Relationship (SAR) study of Lan-P49.1 versus Lan-P49.2. **(A)** Illustration of Lan-P49.1, variant SAR5 and Lan-P49.2. Dashed box indicates the hinge region from Lan-P49.1 swapped into the Lan-P49.2 backbone that created SAR5. Green shading indicates residues different from P49.2. Five rings in the structure are denoted on the top. Residue numbers and hinge region are denoted at the bottom. **(B-N)** Minimal Inhibitory Concentration (MIC) of the lantibiotic variants against human pathogens **(B-F)** and human gut commensals **(G-N)**. Each dot indicates one measurement of MIC value. Each bar indicates median MIC value. Upper and lower dashed lines indicate upper and lower limit of concentration tested, respectively. Observed and expected masses for each compound are listed in Table S1.

## Discussion

Nisin is a class I lantibiotic that has been widely used in the food industry for decades and has been explored as an antimicrobial therapeutic.^22, 25^ Nisin A is produced by *Lactococcus lactis* found in milk. However, how nisin and nisin-like lantibiotics might impact the human gut microbiome has not been elucidated, nor has the presence of nisin-like compounds in the human gut microbiome been investigated in detail. The recent discovery of blauticin and nisin O clearly indicates that nisin-like molecules are encoded in the human microbiome. To first explore the biosynthetic potential of the gut microbiome for lantibiotic production, we systematically performed bioinformatic mining with RODEO for novel nisin-like lantibiotics derived from gut bacterial genomes. Subsequent filtering, multiple sequence alignment and manual curation discovered six gut-derived nisin-like lantibiotics, four of which were not reported previously. Our work therefore greatly expands the repertoire of naturally encoded nisin-like lantibiotics from the human gut.

Utilizing an improved heterologous expression platform for lantibiotics in *E. coli*, we produced all six gut-derived lantibiotics in mixed dehydration states. Blauticin is also in very similar mixed dehydration states when produced by its native host *B. producta*, and its fully dehydrated form has similar efficacy against VRE compared to its mixture of different dehydration states. This finding, along with the previous demonstration that nisin is also produced in various dehydration states in its native producer,^36^ suggests that nisin-like lantibiotics do not require full dehydration for their antimicrobial activity. Therefore, we used purified lantibiotics in mixed dehydration states in this study because it best represents the naturally produced compounds.

Gram-positive pathogens and human gut commensals have varied susceptibility profiles towards the panel of lantibiotics tested. Many mechanisms of lantibiotic resistance are reported in bacteria; besides mechanisms already discussed, an array of two-component systems and cell membrane modification genes have also been reported to confer lantibiotic resistance (see detailed discussion in review ^29^). Besides the presence and number of resistance genes in the genome, the regulation and expression of these genes are also important in lantibiotic resistance. While we did not find strong correlations between the numbers of resistance genes and the MICs of the lantibiotic panel tested, future studies of expression and regulation of various resistance genes in bacteria, especially in human gut commensals, will be needed to elucidate the apparent differences of susceptibility among Gram-positive bacteria. Alternatively, other differences between the tested bacteria such as membrane composition may account for the different susceptibilities.

To further understand the contribution of specific residues in naturally occurring nisin-like lantibiotics to their antimicrobial activity, we studied the SAR of two pairs of closely related lantibiotics: blauticin versus nisin O, and Lan-P49.1 versus Lan-P49.2. Both SARs revealed that the hinge region is critical in determining the efficacy of these lantibiotics, consistent with previous reports.^57^ SAR studies of blauticin versus nisin O also found the first two residues to be important for activity. Surprisingly, when residues in blauticin were changed to their counterparts in nisin O (SAR2 and SAR4), it caused deleterious effects on the antimicrobial activity even against bacteria towards which nisin O has stronger activity than blauticin. Our data suggest that the impact of certain individual residues is context-dependent, implicating intricate interactions among residues along the full length of nisin-like lantibiotics. The 1:1 structure of nisin bound to lipid II has been determined by NMR spectroscopy in DMSO, but the structure of the 2:1 nisin to lipid II complex that is believed to be present in the pores that are made up of eight nisin and four lipid II molecules is currently still not known. Our results showing high sensitivity to replacing even single residues suggest that it may be that the naturally occurring sequences of nisin-like lantibiotics may have evolved such that co-variance of residues is important for forming the 2:1 ratio structure.

In summary, we have discovered gut-derived novel class I lantibiotics through bioinformatic mining, produced them by using an improved heterologous expression platform, and studied their antimicrobial activities against both pathogens and human gut commensals. These characterizations and subsequent SAR studies have revealed the antimicrobial spectrum of both pathogens and human gut commensals, providing insights that will be valuable for future development of lantibiotic-based therapeutics and food preservatives.

## Notes

### Competing Interest Statement

The authors have declared no competing interest.

