## Supporting Information for "Structure-Activity Relationship Studies of Novel Gut-derived Lantibiotics Against Human Gut Commensals"

### Methods

#### General methods

Chemical reagents and media components used in this study were purchased from Sigma-Aldrich or Thermo Fisher Scientific, unless otherwise specified. Oligonucleotides and enzymes were purchased from Integrated DNA Technologies (IDT) and New England Biolabs (NEB), respectively. Polymerase chain reaction (PCR) amplifications were carried out using Q5 polymerase (NEB) on an automated thermocycler (C1000, Bio-Rad). DNA sequencing was performed using appropriate primers by ACGT, Inc. MALDI-TOF MS analyses were conducted at the Mass Spectrometry Facility at UIUC using a Bruker UltrafleXtreme MALDI TOF/TOF spectrometer (Bruker Daltonics). For MALDI−TOF MS analysis, samples were desalted using ZipTipC18 (Millipore) and spotted onto a MALDI target plate with a matrix solution usually consisting of a saturated aqueous solution of super DHB (2,5-dihydroxy benzoic acid; Sigma-Aldrich). Peptide purification was carried out by reversed-phase high performance liquid chromatography (RP−HPLC) on an Agilent 1260 Infinity II instrument equipped with a Macherey-Nagel C18 reverse-phase column (4.6 mm i.d. × 250 mm L). For RP−HPLC, solvent A was 0.1% TFA in $H_2O$, and solvent B was acetonitrile containing 0.1% trifluoroacetic acid (TFA). An elution gradient from 0% solvent B to 100% solvent B over 30 min was used unless specified otherwise.

#### Bioinformatic mining of gut-derived lantibiotic

Bioinformatic mining of gut-derived class I lantibiotics was performed similarly to previously reported.[1] The non-redundant protein records for bacteria in the NCBI RefSeq collection (February 2020) were searched with the LanC-Like Pfam HMM (PF05147.12) using HMMER3 with default settings. Proteins encoded within seven ORFs upstream and downstream of the LanC-Like proteins were searched for precursor peptides using RODEO.[2] These precursor peptides were aligned against both nisin A and blauticin by BLAST, and candidates with less than 30% identity were filtered out. Multiple sequence alignments were performed on nisin A, blauticin and remaining candidate peptides, and close homologs were manually selected by using the following criteria: (a) the peptide length is less than or equal to 72; (b) they contain 5 ring-forming cysteine residues at similar relative locations to nisin A and blauticin. Sequences with different leader peptides but same core peptides were further consolidated.

**Plasmid construction**

The plasmids constructed for this study and their combined application for peptide production and purification are shown in Figure S2. The *Blautia producta SCSK* ($BP_{SCSK}$) $tRNA^{Glu}$ sequences were identified using the algorithm tRNAscan-SE.[3, 4] For blauticin plasmid construction, pDuet1-His-BpcA and pCDFDuet1-BpcBC were synthesized by SynbioTechnologies. The genes $BP_{SCSK}$ $tRNA^{Glu}$ and gluRS were constructed using synthetic gene fragment gBlocks (IDT), and amplified using primers for both inserts and backbones (Table S2). For blauticin analog or SAR mutant precursor peptide plasmid construction, the genes for precursors were constructed using synthetic gene fragment gBlocks (IDT), and amplified using primers for both inserts and backbones (Table S2).

**Heterologous expression of peptides with pEVOL plasmids**

Chemically competent *E. coli* BL21 (DE3) cells were transformed with combinations of the plasmids prepared (Figure S2) to co-express enzymes. Colonies were selected on LB agar plates containing the appropriate antibiotics listed for each plasmid. Single colonies were used to inoculate 5 mL of overnight starter cultures in LB with the appropriate antibiotic. The starter cultures were then used to inoculate 1 L of TB cultures containing the appropriate antibiotic, which were incubated at 37 °C with shaking (180 rpm) until they reached the beginning of the exponential growth phase (an $OD_{600}$ of approximately 0.4). Cultures were then cooled at 4 °C for 15 min prior to the addition of L-arabinose to a final concentration of 0.1%. Cultures were placed into a 20 °C incubator and shaken (180 rpm) for at least 2 h until the culture reached the end of the exponential growth phase (an $OD_{600}$ of approximately 1.2). The "slow growing time" that was maintained for a minimal of two hours before adding the next inducer was critical for the successful expression. Next, isopropyl β-D-1-thiogalactopyranoside (IPTG) was added to the culture to a final concentration of 0.5 mM. Cultures were placed into an 18 °C incubator and shaken overnight (180 rpm). Cells were harvested by centrifugation at 4,500 × g for 20 min. The culture media was decanted, and the cell pellet stored at −80 °C prior to purification.

**Peptide purification**

Cell pellet was suspended in 30 mL of lysis buffer (20 mM $NaH_2PO_4$, 500 mM NaCl, 0.5 mM imidazole, 20% glycerol, pH 7.5 at 25 °C) supplemented with Pierce Protease Inhibitor tablets (1 tablet per 1 L culture; Fisher Scientific) and benzonase (1 U for every 20 mL of lysate). The cell pellet was lysed by sonication (50% amplitude, 2 s pulse, 5 s pause, 15 min). Crude lysate was centrifuged at 24,000 ×g for 60 min at 4 °C and the supernatant was obtained. The supernatant was loaded to onto a His60 Ni superflow resin (Takara) equilibrated with lysis buffer, and cell lysate was applied three times to it under the action of gravity. After loading the sample, the resin was washed with two column volumes (CV) of washing buffer (4 M guanidine hydrochloride, 20 mM $NaH_2PO_4$, 300 mM NaCl, 30 mM imidazole, pH 7.5 at 25 °C), and eluted using 1 CV of LanA Elution Buffer (4 M guanidine hydrochloride, 20 mM Tris HCl, pH 7.5 at 25 °C, 100 mM NaCl, 1 M imidazole). Peptides eluted from the Ni resin were concentrated and desalted using 3 kDa Amicon filters (Millipore). For leader peptide removal, the peptide (1:20 trypsin to peptide) was incubated with the trypsin protease in 50 mM Tris-HCl, pH 8 overnight at 37 °C. Lastly, the core peptides were further purified with HPLC using an Agilent 1200 instrument (Agilent) as specified in general methods.

**Purification of blauticin from *Blautia producta* SCSK**

Stationary anaerobic culture of $BP_{SCSK}$ was inoculated 1:100 to 1 L BHIS. Culture was incubated on a heating plate at 37 °C with 220 rpm stirring for 2 days in an anaerobic chamber. Bacteria were pelleted with 15,000 x g centrifugation for 15 min. The supernatant was acidified with acetic acid to pH 5.5 and filtered through a 0.22 um filter. Filtered supernatant was loaded onto a HiTrap SP HP column using an AKTA Pure FPLC, which was eluted sequentially with 3 CV of distilled water, 200 mM NaCl, 400 mM NaCl, 600 mM NaCl, 800 mM NaCl, and 1 M NaCl. The 800 mM NaCl eluate was concentrated with a 3 kDa Amicon filter, followed by dialyzation with DPBS using a 3 kDa Amicon filter. Purification of blauticin was confirmed by SDS-PAGE, VRE inhibition in vitro assay, and MALDI-TOF MS.

**Isolation of human gut commensal bacteria**

Isolation and growth of commensal bacteria was done as previously reported.[5] Briefly, fresh donor fecal samples were transferred into an anaerobic chamber (Coy Labs) within 1 h of collection. The anaerobic chamber was maintained using a gas mix consisting of 5% hydrogen, 5% carbon dioxide and 90% nitrogen, and hydrogen was maintained at ~3.0% using an anaerobic gas infuser. Fecal samples were resuspended in pre-reduced PBS and plated on Columbia blood agar or brain-heart infusion (BHI – Difco, BD) agar plates in three serial 10-fold dilutions and incubated at 37 ºC for 48 – 96 h. Isolated colonies were re-streaked for purity onto Columbia blood agar and frozen in pre-reduced 10% glycerol in PBS.

**Bacterial growth**

For broth cultures, isolates were grown in pre-reduced BHI supplemented with 5 g/L yeast extract (Difco, BD) and 0.1% L-cysteine under anaerobic condition.

**Minimal Inhibition Concentration (MIC) assay**

All MIC assays were performed under anaerobic condition. Stationary bacterial culture in BHIS (BHI supplemented with 5 g/L yeast extract and 0.1% L-cysteine) were inoculated 1:100 to wells containing 200 μL BHIS in U-bottom 96-well plates. Lantibiotics were dissolved in DPBS, and concentrations were determined by BCA assay (Pierce) according to the manufacturer's protocol. Lantibiotic stock solutions were added to wells to reach desired final concentration, with 3-fold serial dilution across rows to create concentration gradients. Bacteria-inoculated wells without lantibiotics served as control. Plates were incubated at 37 ºC for 48 h before being taken out of the anaerobic chamber for centrifugation. Plates were imaged with an iBright 1500 imager (Invitrogen). The lowest concentration of lantibiotics that caused no or significantly less bacterial pellets than those without lantibiotics were identified as MIC.

**Lantibiotic resistance gene mining**

Genomes of pathogens and human gut commensals investigated were retrieved from RefSeq (Table S3). Genomes were annotated using Prokka,[6] which is based on both Hidden Markov Model (HMM) search and BLAST search. Genes with consistent annotations were numerated in each genome. For lantibiotic resistance genes without consistent annotations, an HMM search with corresponding HMM profiles (Table S3) were performed with hmmsearch in HMMER 3.3.[7] In case of *mprF* and *nsr* genes where HMM profiles were not available, individual protein sequences were retrieved from Uniprot (Table S3) and were searched against genomes with phmmer in HMM 3.3.[7] *E*-value cutoffs were set at $e^{-15}$ for both hmmsearch and phmmer.

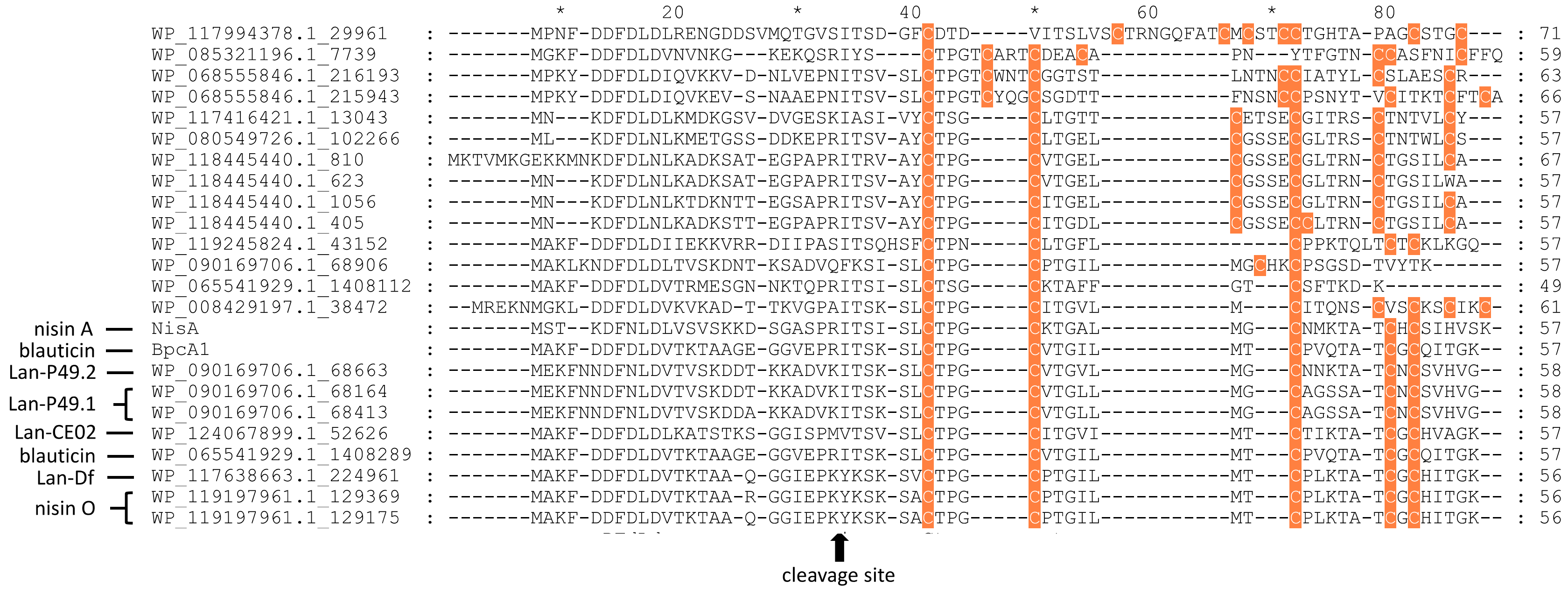


**Fig. S1 Multiple Sequence Alignment (MSA) of nisin-like lantibiotic candidates discovered in the RefSeq database after length filtering.** Cysteine residues that are important for thioether ring formation are colored in orange. The precursor peptides for nisin A (NisA) and blauticin (BpcA1) were included for comparison of cysteine positions as benchmarks for manual curation. The sequence length of each peptide is shown on the far right. Protease cleavage sites of precursor peptides of nisin A and blauticin are marked at the bottom. Final nisin-like lantibiotic candidates are denoted on the left.

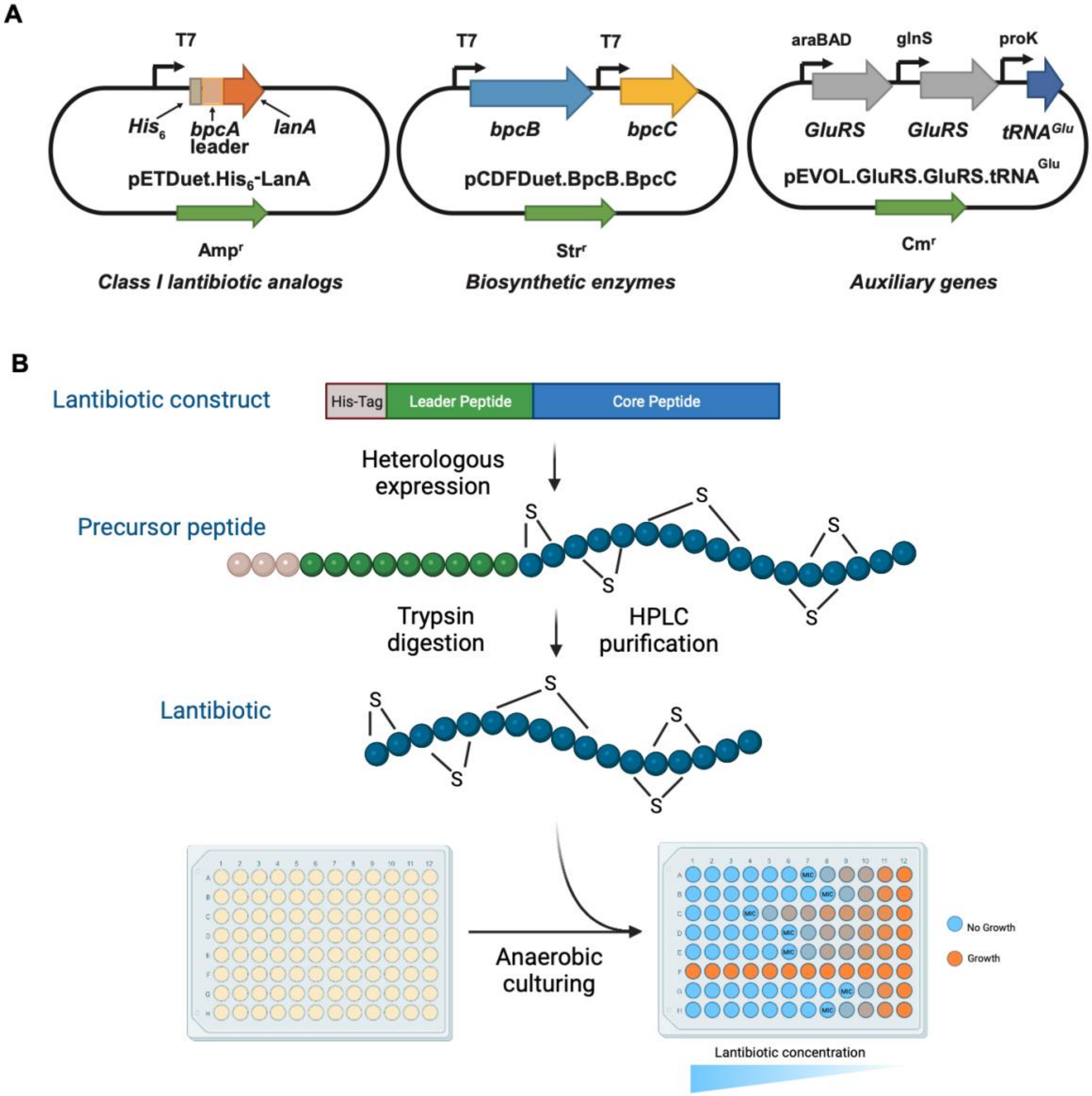


**Fig. S2 Scheme of improved expression platform and lantibiotic production workflow. (A)** Illustration of the three plasmids used in the expression platform. **(B)** Workflow of expression, purification, and MIC testing of each lantibiotic variant. Figure was created with biorender.com.

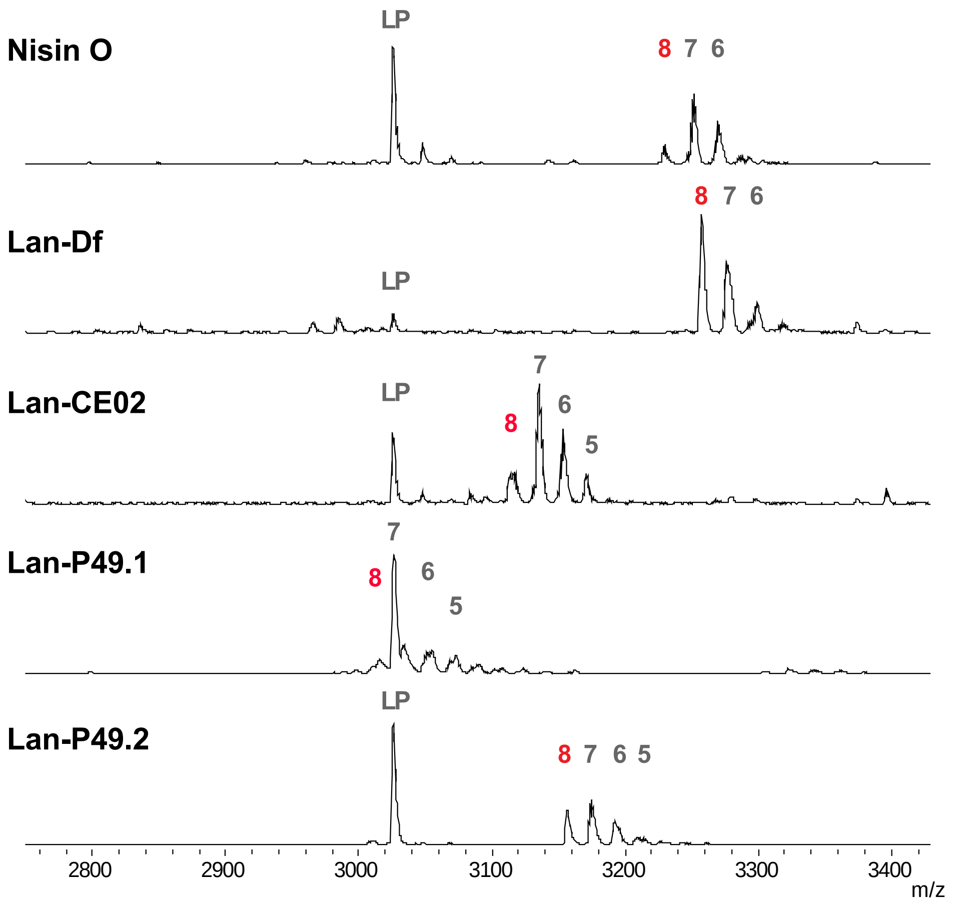


**Fig. S3 MALDI-TOF MS analysis of nisin O, Lan-Df, Lan-CE02, Lan-P49.1, and Lan-P49.2 produced in *E. coli*.** The numbers on top of the peaks indicate dehydration states for each peptide. Red numbers indicate full dehydration of all Ser/Thr residues. LP, leader peptide. Observed and expected masses for each compound are listed in Table S1. For Lan-P49.1, the peaks of the seven-fold dehydrated peptide and the leader peptide overlap.

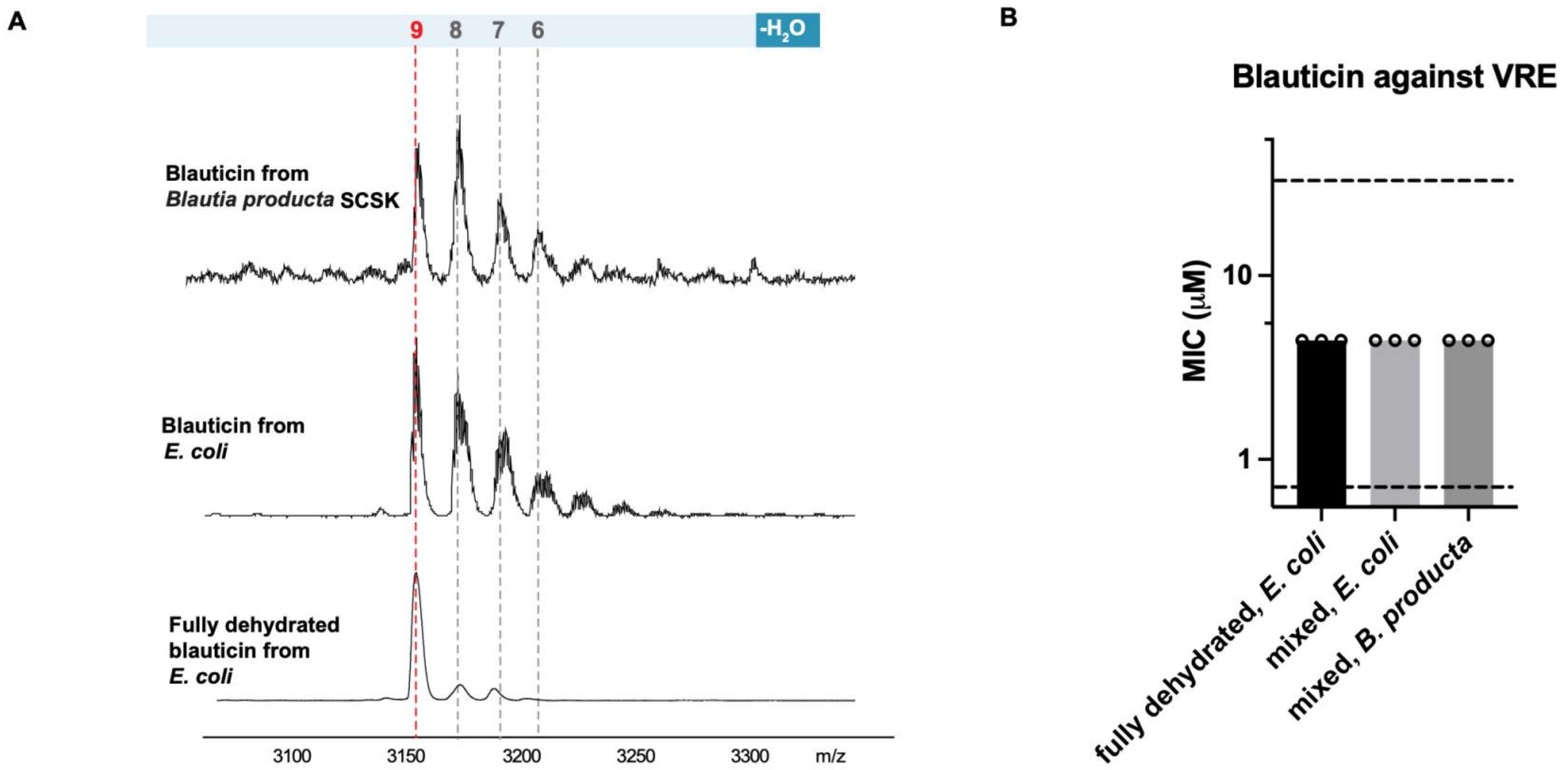


**Fig. S4 Bioactivities of blauticin heterologously produced in *E. coli* and purified from its native host *B. producta*.** MIC assay of blauticin in fully dehydrated form or mixed forms purified from *E. coli*, and blauticin in mixed forms from *B. producta* against Vancomycin-resistant *Enterococcus faecium* ATCC700221 (VRE). Each dot indicates one measurement of MIC value. Each bar indicates median MIC value. Upper and lower dashed lines indicate upper and lower limit of concentration tested, respectively.

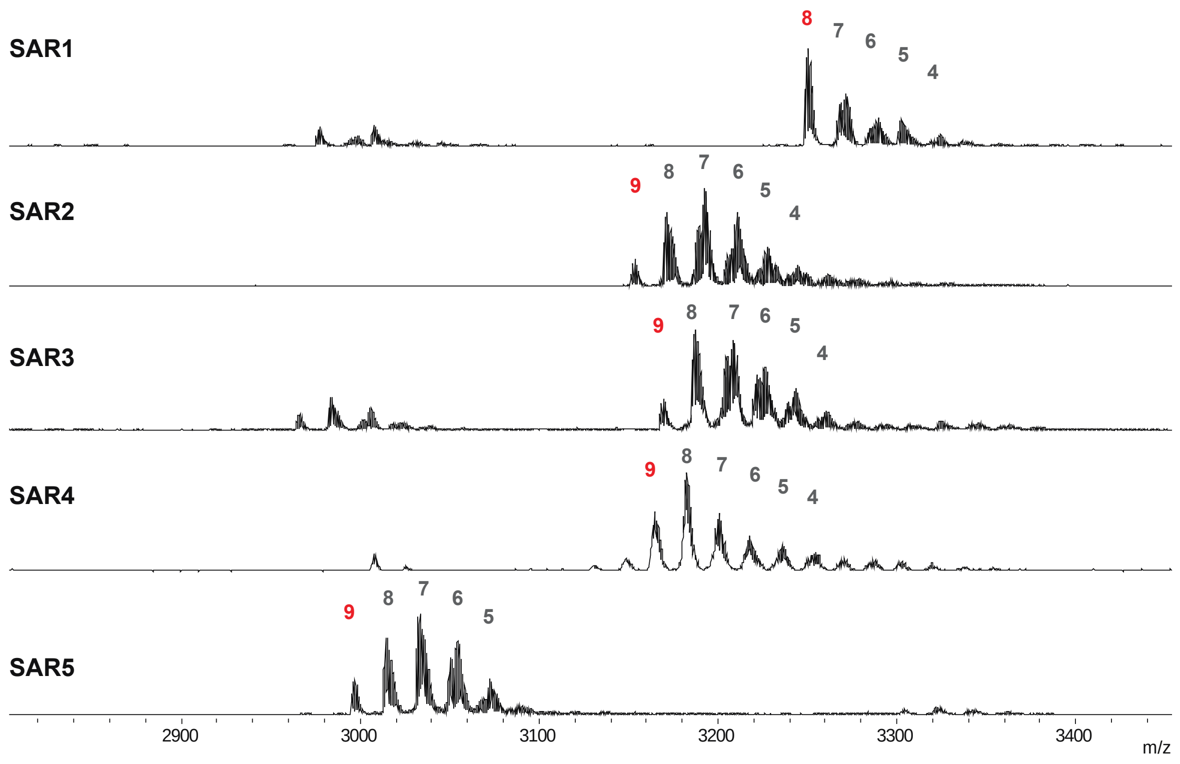


**Fig. S5 MALDI-TOF mass spectra of purified blauticin variants.** The numbers on top of the peaks indicate dehydration states for each peptide. Red numbers indicate fully dehydrated forms. Observed and expected masses for each compound are listed in Table S1.

**Table S1. Lanthipeptide expected and observed masses (Da).***

| Number of dehydrations | 0 | 1 | 2 | 3 | 4 | 5 | 6 | 7 | 8 | 9 |
|---|---|---|---|---|---|---|---|---|---|---|
| **Blauticin (expected)** | 3316 | 3298 | 3280 | 3262 | 3244 | 3226 | 3208 | 3190 | 3172 | 3154 |
| **Blauticin (observed)** | NO | NO | NO | NO | 3245 | 3228 | 3210 | 3192 | 3175 | 3156 |
| **Nisin O (expected)** | 3373 | 3355 | 3337 | 3319 | 3301 | 3283 | 3265 | 3247 | 3229 | NA |
| **Nisin O (observed)** | NO | NO | NO | NO | NO | 3283 | 3267 | 3248 | 3228 | NA |
| **Lan-Df (expected)** | 3401 | 3383 | 3365 | 3347 | 3329 | 3311 | 3293 | 3275 | 3257 | NA |
| **Lan-Df (observed)** | NO | NO | NO | NO | NO | NO | 3294 | 3275 | 3257 | NA |
| **Lan-CE02 (expected)** | 3257 | 3239 | 3221 | 3203 | 3185 | 3167 | 3149 | 3131 | 3113 | NA |
| **Lan-CE02 (observed)** | NO | NO | NO | NO | NO | 3171 | 3151 | 3133 | 3113 | NA |
| **Lan-P49.1 (expected)** | 3159 | 3141 | 3123 | 3105 | 3087 | 3069 | 3051 | 3033 | 3015 | NA |
| **Lan-P49.1 (observed)** | NO | NO | NO | 3105 | 3087 | 3069 | 3051 | 3028 | 3014 | NA |
| **Lan-P49.2 (expected)** | 3300 | 3282 | 3264 | 3246 | 3228 | 3210 | 3192 | 3174 | 3156 | NA |
| **Lan-P49.2 (observed)** | NO | NO | NO | NO | NO | 3210 | 3193 | 3173 | 3155 | NA |
| **SAR1 (expected)** | 3393 | 3375 | 3357 | 3339 | 3321 | 3303 | 3285 | 3267 | 3249 | NA |
| **SAR1 (observed)** | NO | NO | NO | 3336 | 3322 | 3303 | 3287 | 3266 | 3248 | NA |
| **SAR2 (expected)** | 3314 | 3296 | 3278 | 3260 | 3242 | 3224 | 3205 | 3187 | 3169 | 3151 |
| **SAR2 (observed)** | NO | NO | NO | 3261 | 3243 | 3225 | 3207 | 3189 | 3170 | 3151 |
| **SAR3 (expected)** | 3330 | 3312 | 3294 | 3276 | 3258 | 3240 | 3222 | 3204 | 3186 | 3168 |
| **SAR3 (observed)** | NO | NO | NO | 3275 | 3259 | 3241 | 3225 | 3105 | 3186 | 3168 |
| **SAR4 (expected)** | 3325 | 3307 | 3289 | 3271 | 3253 | 3235 | 3217 | 3198 | 3180 | 3162 |
| **SAR4 (observed)** | NO | 3305 | 3287 | 3270 | 3251 | 3235 | 3217 | 3198 | 3180 | 3163 |
| **SAR5 (expected)** | 3157 | 3139 | 3121 | 3103 | 3085 | 3067 | 3049 | 3031 | 3013 | 2995 |
| **SAR5 (observed)** | NO | NO | NO | NO | NO | 3069 | 3051 | 3032 | 3013 | 2996 |

($^{*}$All masses are in Dalton; NA = not applicable; NO = not observed).

**Table S2. List of oligonucleotides and gene fragments used in this study.**

| Name | **5’ Sequence 3’** |
|---|---|
| pEVOL_GluRS1 (BPscsk)_gBlock | CCCGTTTTTTTGGGCTAACAGGAGGAATTAGATCTATGTCCACAGTGCGCACCCGTTTTGCTCCGTCTCCGACGGGTCG<br>TATGCATGTAGGAAATCTGCGTACCGCACTGTACGCATATTTAATTGCCAAACACGAAAACGGCTCATTCATGCTGCGT<br>ATTGAAGATACTGATCAAGAACGTTTTGTAGACGGTGCCCTGGAAATCATTTATCGCACGTTAGCCAAGACGGGCTTGA<br>TTCATGACGAGGGCCCGGATAAAGATGGAGGCTATGGTCCGTATGTACAGAGCGAACGTAATGCACAAGGCATTTATCT<br>CAACTACGCCAAACAACTGATTGAACAGGGTGATGCCTACTATTGCTTTTGTACGAAGGAACGGTTAGATTCCCTGAAG<br>GCGTCTGTAGGAGAAGATGGTAAAGAAATTGCTGTGTATGATAAACACTGCTTACATCTTTCTCGTGAGGAGGTGGAGG<br>CGAAACTGGCGGCCGGCGAACCGCATGTGATCCGTTTTAATATGCCAACGGAAGGGAACACCACCTTTCATGATGATAT<br>CTACGGAGACATCACGGTGCCGAATAATGAGCTGGACGACCTTATTCTGATCAAATCGGATGGTTACCCGACTTATAAC<br>TTTGCTAACGTCATTGATGATCACTTGATGGGTATCACGCACGTTGTGCGCGGCAATGAATATCTTAGCTCTAGCCCAA<br>AATATAATCGGATTTACGAAGCCTTTGGTTGGGAAATCCCGACATATGTGCACTGCCCGTTGATTACCAACGAAGAGCA<br>TAAGAAACTGTCCAAGCGTAGCGGGCACTCATCTTATGAAGACTTAACGGACCAAGGCTTCCTCACCGAAGCCATCGTT<br>AATTACGTAGCTCTGCTCGGGTGGTGCCCGGAGGATAATCGCGAGATCTTCAGTCTGGAAGAACTGGTGAAAGAGTTTG<br>ATTATCACCATATGAGTAAATCCCCGGCCGTCTTTGATATGACCAAGCTGAAGTGGATGAACGGGGAATATATTAAAGC<br>CATGGATTTTGATAAATTTTGTGATATGGCCCTGCCTTTCGTGAAAGAAGCGGTTAAAAGTGATCTGGATCTGAAAAAA<br>ATCGTCCAGATGGTGAAAACGCGCATTGAAGTCTTCCCCGATATTCCAGCCCTTATCGATTTCTTTGAAGAAGTGCCGG<br>AATACGATGTTTCTATGTATACCCATAAAAAAATGAAAACTAACCCGGAACTCTCCTTGGAGGTTCTGAAAAAAATCCT<br>GCCAGTACTGGAAGATTTTGAGGACTATACCAATGATGCTTTATATGATTTACTGTGTGGCTTTGCGAAAGAGAACGGC<br>TATAAGAACGGTCAGATCCTGTGGCCGATTCGTACAGCGCTGAGCGGGAAGCAGATGACTCCAGCGGGAGCTACCGAGA<br>TCTTGGAGGTGCTGGGTAAAGAAGAATCGATGAAGCGTCTTCATGCGGCCGTGGAAAAACTGGAATGCGTTTGAGTCGA<br>CCATCATCATCATCATCATTGAGTTTAAACGGTCTCCAGCTTGGCTGTTTTGGCGGATGAGAGAAGATTTTCAGCCTGA<br>TACAGATTAAATCAGAACGCAGAAGCGGTCTGATAAAACAGAATTTGCCTGGCGGCAGTAGCGCGGTGGTCCCACCTGA<br>CCCCATGCCGAACTCAGAAGTGAAACGCCGTAGCGCCGATGG |
| pEVOL_GluRS2 (BPscsk)_gBlock | CAGAAGTGAAACGCCGTAGCGCCGATGGTAGTGTGGGGTCTCCCCATGCGAGAGTAGGGAACTGCCAGGCATCAAATAA<br>AACGAAAGGCTCAGTCGAAAGACTGGGCCTTGTTTGTGAGCTCCCGGTCATCAATCATCCCCATAATCCTTGTTAGATT<br>ATCAATTTTAAAAAACTAACAGTTGTCAGCCTGTCCCGCTTTAATATCATACGCCGTTATACGTTGTTTACGCTTTGAG<br>GAATCCCATATGTCCACAGTGCGCACCCGTTTTGCTCCGTCTCCGACGGGTCGTATGCATGTAGGAAATCTGCGTACCG<br>CACTGTACGCATATTTAATTGCCAAACACGAAAACGGCTCATTCATGCTGCGTATTGAAGATACTGATCAAGAACGTTT<br>TGTAGACGGTGCCCTGGAAATCATTTATCGCACGTTAGCCAAGACGGGCTTGATTCATGACGAGGGCCCGGATAAAGAT<br>GGAGGCTATGGTCCGTATGTACAGAGCGAACGTAATGCACAAGGCATTTATCTCAACTACGCCAAACAACTGATTGAAC<br>AGGGTGATGCCTACTATTGCTTTTGTACGAAGGAACGGTTAGATTCCCTGAAGGCGTCTGTAGGAGAAGATGGTAAAGA<br>AATTGCTGTGTATGATAAACACTGCTTACATCTTTCTCGTGAGGAGGTGGAGGCGAAACTGGCGGCCGGCGAACCGCAT<br>GTGATCCGTTTTAATATGCCAACGGAAGGGAACACCACCTTTCATGATGATATCTACGGAGACATCACGGTGCCGAATA<br>ATGAGCTGGACGACCTTATTCTGATCAAATCGGATGGTTACCCGACTTATAACTTTGCTAACGTCATTGATGATCACTT<br>GATGGGTATCACGCACGTTGTGCGCGGCAATGAATATCTTAGCTCTAGCCCAAAATATAATCGGATTTACGAAGCCTTT |

| | |
|---|---|
| | GGTTGGGAAATCCCGACATATGTGCACTGCCCGTTGATTACCAACGAAGAGCATAAGAAACTGTCCAAGCGTAGCGGGC<br>ACTCATCTTATGAAGACTTAACGGACCAAGGCTTCCTCACCGAAGCCATCGTTAATTACGTAGCTCTGCTCGGGTGGTG<br>CCCGGAGGATAATCGCGAGATCTTCAGTCTGGAAGAACTGGTGAAAGAGTTTGATTATCACCATATGAGTAAATCCCCG<br>GCCGTCTTTGATATGACCAAGCTGAAGTGGATGAACGGGGAATATATTAAAGCCATGGATTTTGATAAATTTTGTGATA<br>TGGCCCTGCCTTTCGTGAAAGAAGCGGTTAAAAGTGATCTGGATCTGAAAAAAATCGTCCAGATGGTGAAAACGCGCAT<br>TGAAGTCTTCCCCGATATTCCAGCCCTTATCGATTTCTTTGAAGAAGTGCCGGAATACGATGTTTCTATGTATACCCAT<br>AAAAAAATGAAAACTAACCCGGAACTCTCCTTGGAGGTTCTGAAAAAAATCCTGCCAGTACTGGAAGATTTTGAGGACT<br>ATACCAATGATGCTTTATATGATTTACTGTGTGGCTTTGCGAAAGAGAACGGCTATAAGAACGGTCAGATCCTGTGGCC<br>GATTCGTACAGCGCTGAGCGGGAAGCAGATGACTCCAGCGGGAGCTACCGAGATCTTGGAGGTGCTGGGTAAAGAAGAA<br>TCGATGAAGCGTCTTCATGCGGCCGTGGAAAAACTGGAATGCGTTTGACTGCAGTTTCAAACGCTAAATTGCCTGATGC<br>GCTAC |
| pEVOL_CUCtRNA(BPscsk)_gBlock | GAATGCGTTTGACTGCAGTTTCAAACGCTAAATTGCCTGATGCGCTACGCTTATCAGGCCTACATGATCTCTGCAATAT<br>ATTGAGTTTGCGTGCTTTTGTAGGCCGGATAAGGCGTTCACGCCGCATCCGGCAAGAAACAGCAAACAATCCAAAACGC<br>CGCGTTCAGCGGCGTTTTTTCTGCTTTTCTTCGCGAATTAATTCCGCTTCGCAACATGTGAGCACCGGTTTATTGACTA<br>CCGGAAGCAGTGTGACCGTGTGCTTCTCAAATGCCTGAGGCCAGTTTGCTCAGGCTCTCCCCGTGGAGGTAATAATTGA<br>CGATATGATCAGTGCACGGCTAACTAAGCGGCCTGCTGACTTTCTCGCCGATCAAAAGGCATTTTGCTATTAAGGGATT<br>GACGAGGGCGTATCTGCGCAGTAAGATGCGCCCCGCATTGGTTCGTTGGTCAAGCGGTtAAGACGCCGCCCTCTCACGG<br>CGGAAaCAGGGGTTCGATTCCCCTACGAACTGCAATTCGAAAAGCCTGCTCAACGAGCAGGCTTTTTTGCATGCTCGAG<br>CAGCTCAGGGTCGAATTTGCTTTCGAATTTCTGCCATTCATCCGCTTATTATCACTTATTCAGGCGTAGCAAC |
| pEVOL_UUCtRNA(BPscsk)_gBlock | GCAGTTTCAAACGCTAAATTGCCTGATGCGCTACGCTTATCAGGCCTACATGATCTCTGCAATATATTGAGTTTGCGTG<br>CTTTTGTAGGCCGGATAAGGCGTTCACGCCGCATCCGGCAAGAAACAGCAAACAATCCAAAACGCCGCGTTCAGCGGCG<br>TTTTTTCTGCTTTTCTTCGCGAATTAATTCCGCTTCGCAACATGTGAGCACCGGTTTATTGACTACCGGAAGCAGTGTG<br>ACCGTGTGCTTCTCAAATGCCTGAGGCCAGTTTGCTCAGGCTCTCCCCGTGGAGGTAATAATTGACGATATGATCAGTG<br>CACGGCTAACTAAGCGGCCTGCTGACTTTCTCGCCGATCAAAAGGCATTTTGCTATTAAGGGATTGACGAGGGCGTATC<br>TGCGCAGTAAGATGCGCCCCGCATTGGCTCCATGGTCAAGCGGTtAAGACACCGCCCTTTCACGGCGGTAaCAGGGGTT<br>CAAATCCCCTTGGAGTCACAATTCGAAAAGCCTGCTCAACGAGCAGGCTTTTTTGCATGCTCGAGCAGCTCAGGGTCGA<br>ATTTGCTTTCGAATTTCTGCCATTCATCCGCTTATTATCACTTATTCAGGCGTAGCAACCAGGCGTTTAAGGGCACCAA<br>TAACTGCCTTAAAAAAATTACGCCCCGCCCTGCCACTCATCGCAGTACTGTTGTAATTCATTAAGCATTCTGCCGACAT<br>GGAAGCCATCACAAACGGCATGATGAACCTGAATC |
| pEVOL_GluRS1(BPscsk)F | CGTTTTTTTGGGCTAACAGGAG |
| pEVOL_GluRS1(BPscsk)R | GCGCTACGGCGTTTC |
| pEVOL_GluRS2(BPscsk)F | CAGAAGTGAAACGCCGTAG |
| pEVOL_GluRS2(BPscsk)R | GTAGCGCATCAGGCAATTTAG |
| pEVOL_BB_F | CTCGAGCAGCTCAGG |
| pEVOL_BB_R | GAGACGGAGCAAAACGGGTGCGCACTGTGGACATAGATCTAATTCCTCCTGTTAGC |

| | |
|---|---|
| pEVOL_CUCtRNA(BPscsk)F | GAATGCGTTTGACTGCAGTTTCAAACGCTAAATTGCCTGA |
| pEVOL_CUCtRNA(BPscsk)R | GTTGCTACGCCTGAATAAGTGATAATAAGCGGATGAATG |
| pEVOL_UUCtRNA(BPscsk)F | GCAGTTTCAAACGCTAAATTG |
| pEVOL_UUCtRNA(BPscsk)R | GATTCAGGTTCATCATGCC |
| **Blauticin Analogs** | |
| pETDuet_nisin O_gBlock | ATAACAATTCCCCTCTAGAAATAATTTTGTTTAACTTTAAGAAGGAGATATACCATGGGCAGCAGCCATCACCATCATCACCACAGCCAGGATCCGAATTCGAGCTCGGCGCGCCTGCAGGTCGACATGGCGAAATTCGACGACTTCGATCTGGACGTTACCAAAACCGCAGCTGGCGAAGGCGGTGTTGAACCTCGTTATAAAGTAAATCTGCTTGTACGCCAGGCTGCCCGACCGGCATTCTGATGACTTGCCCACTGAAAACGGCAACCTGTGGATGTCATATTACCGGCAAATAA |
| pETDuet_D. formicigenerans_gBlock | ATAACAATTCCCCTCTAGAAATAATTTTGTTTAACTTTAAGAAGGAGATATACCATGGGCAGCAGCCATCACCATCATCACCACAGCCAGGATCCGAATTCGAGCTCGGCGCGCCTGCAGGTCGACATGGCGAAATTCGACGACTTCGATCTGGACGTTACCAAAACCGCAGCTGGCGAAGGCGGTGTTGAACCGCGTTATAAATCCAAAAGCGTGTGCACCCCGGGCTGTCCGACAGGTATTCTGATGACCTGCCCGCTGAAAACCGCCACCTGTGGGTGTCATATTACCGGCAAGTAA |
| pETDuet_Clostridum sp E02_gBlock | ATAACAATTCCCCTCTAGAAATAATTTTGTTTAACTTTAAGAAGGAGATATACCATGGGCAGCAGCCATCACCATCATCACCACAGCCAGGATCCGAATTCGAGCTCGGCGCGCCTGCAGGTCGACATGGCGAAATTCGACGACTTCGATCTGGACGTTACCAAAACCGCAGCTGGCGAAGGCGGTGTTGAACCGCGTGTGACGTCCGTGTCCTTATGTACGCCGGGATGCATCACGGGCGTAATCATGACTTGTACCATTAAAACGGCAACCTGTGGCTGCCACGTGGCCGGCAAATAA |
| pETDuet_Pseudobutyrivibrio_sp49_1_gBlock | ATAACAATTCCCCTCTAGAAATAATTTTGTTTAACTTTAAGAAGGAGATATACCATGGGCAGCAGCCATCACCATCATCACCACAGCCAGGATCCGAATTCGAGCTCGGCGCGCCTGCAGGTCGACATGGCGAAATTCGACGACTTCGATCTGGACGTTACCAAAACCGCAGCTGGCGAAGGCGGTGTTGAACCGCGCATCACTTCGAAATCTCTGTGTACCCCGGGTTGCGTCACCGGGCTGCTTATGGGTTGTGCCGGCAGCTCTGCCACATGTAATTGTAGTGTGCATGTTGGCTAA |
| pETDuet_Pseudobutyrivibrio_sp49_2_gBlock | ATAACAATTCCCCTCTAGAAATAATTTTGTTTAACTTTAAGAAGGAGATATACCATGGGCAGCAGCCATCACCATCATCACCACAGCCAGGATCCGAATTCGAGCTCGGCGCGCCTGCAGGTCGACATGGCGAAATTCGACGACTTCGATCTGGACGTTACCAAAACCGCAGCTGGCGAAGGCGGTGTTGAACCGCGCATCACATCGAAATCGCTGTGCACTCCAGGTTGCGTGACCGGTGTACTGATGGGGTGTAACAACAAAACCGCAACTTGTAATTGCAGTGTCCACGTCGGATGA |
| pETDuet_BB_F | CATAATGCTTAAGTCGAACAGAAAG |
| pETDuet_BB_R | CAAAATTATTTCTAGAGGGGAATTGTTAT |
| pETDuet_nisin_DF_CE02_sp49_F | GAATTGTGAGCGGATAACAATTCCCCTCTAGAAATAATTTTG |
| pETDuet_nisin O_R | TGTTCGACTTAAGCATTATGCGGCCGCTTATTTGCCGGTAATATGACATCC |

| | |
|---|---|
| pETDuet_D. formicigenerans_R | TGTTCGACTTAAGCATTATGCGGCCGCTTACTTGCCGGTAATATGACAC |
| pETDuet_Clostridum sp E02_R | TGTTCGACTTAAGCATTATGCGGCCGCTTATTTGCCGGCCACG |
| pETDuet_Pseudobutyrivibrio_sp49_1_R | TGTTCGACTTAAGCATTATGCGGCCGCTTAGCCAACATGCACACTAC |
| pETDuet_Pseudobutyrivibrio_sp49_2_R | TGTTCGACTTAAGCATTATGCGGCCGCTCATCCGACGTGGAC |
| **Blauticin SAR** | |
| pETDuet_SAR1_ITtoYK_gBlock | ATGGGCAGCAGCCATCACCATCATCACCACAGCCAGGATCCGAATTCGAGCTCGGCGCGCCTGCAGGTCGACATGGCGAAATTCGACGACTTCGATCTGGACGTTACCAAAACCGCAGCTGGCGAAGGCGGTGTTGAACCGCGTTATAAATCCAAAAGCCTGTGTACCCCGGGTTGCGTTACCGGTATTCTGATGACCTGTCCGGTTCAAACCGCTACCTGTGGTTGTCAGATCACCGGCAAATAA |
| pETDuet_SAR2_V12P_gBlock | ATGGGCAGCAGCCATCACCATCATCACCACAGCCAGGATCCGAATTCGAGCTCGGCGCGCCTGCAGGTCGACATGGCGAAATTCGACGACTTCGATCTGGACGTTACCAAAACCGCAGCTGGCGAAGGCGGTGTTGAACCGCGTATTACCTCCAAAAGCCTGTGTACCCCGGGTTGCCCGACCGGTATTCTGATGACCTGTCCGGTTCAAACCGCTACCTGTGGTTGTCAGATCACCGGCAAATAA |
| pETDuet_SAR3_VQtoLK_gBlock | ATGGGCAGCAGCCATCACCATCATCACCACAGCCAGGATCCGAATTCGAGCTCGGCGCGCCTGCAGGTCGACATGGCGAAATTCGACGACTTCGATCTGGACGTTACCAAAACCGCAGCTGGCGAAGGCGGTGTTGAACCGCGTATTACCTCCAAAAGCCTGTGTACCCCGGGTTGCGTTACCGGTATTCTGATGACCTGTCCGCTGAAAACCGCTACCTGTGGTTGTCAGATCACCGGCAAATAA |
| pETDuet_SAR4_Q29H_gBlock | ATGGGCAGCAGCCATCACCATCATCACCACAGCCAGGATCCGAATTCGAGCTCGGCGCGCCTGCAGGTCGACATGGCGAAATTCGACGACTTCGATCTGGACGTTACCAAAACCGCAGCTGGCGAAGGCGGTGTTGAACCGCGTATTACCTCCAAAAGCCTGTGTACCCCGGGTTGCGTTACCGGTATTCTGATGACCTGTCCGGTTCAAACCGCTACCTGTGGTTGTCATATCACCGGCAAATAA |
| pETDuet_SAR5_sp49_2_AGS_gBlock | ATGGGCAGCAGCCATCACCATCATCACCACAGCCAGGATCCGAATTCGAGCTCGGCGCGCCTGCAGGTCGACATGGCGAAATTCGACGACTTCGATCTGGACGTTACCAAAACCGCAGCTGGCGAAGGCGGTGTTGAACCGCGTATCACATCGAAATCGCTGTGCACTCCAGGTTGCGTGACCGGTGTACTGATGGGGTGTGCCGGCAGCACCGCAACTTGTAATTGCAGTGTCCACGTCGGATGA |
| pETDuerSAR_BB_F | CCGCATAATGCTTAAGT |
| pETDuerSAR_BB_R | CCATGGTATATCTCCTTC |
| pETDuerSAR_SAR12345_F | GAAGGAGATATACCATGGGCAGCAGC |
| pETDuerSAR_SAR123_R | CATTATGCGGCCGCTTATTTGCCGGTGATCTG |

| | |
|---|---|
| pETDuerSAR_SAR4_R | CATTATGCGGCCGCTTATTTGCCGGTGATATGAC |
| pETDuerSAR_SAR5_R | CATTATGCGGCCGCTCATCCGACGTGGAC |

**Table S3. List of bacterial genomes and PFAM HMM used in this study.**

| **Pathogens** | **NCBI Accession** |
|---|---|
| Enterococcus faecium ATCC700221 | GCF_009734005.1 |
| Staphylococcus aureus USA300 | GCF_017834975.1 |
| Staphylococcus epidermidis SK135 | GCF_000177115.1 |
| Listeria monocytogenes 10403S | GCF_000168695.2 |
| Clostridioides difficile VPI10463 | GCA_001995155.1 |
| **Commensals** | **NCBI Accession** |
| Anaerostipes hadrus MSK.14.23 | GCF_013302595.1 |
| Blautia obeum MSK.18.40 | GCF_013299585.1 |
| Blautia luti MSK.20.18 | GCF_020553125.1 |
| Erysipeloclostridium ramosum MSK.23.96 | GCF_019125475.1 |
| Dorea formicigenerans MSK.17.61 | GCF_013300535.1 |
| Coprococcus comes MSK.11.23 | GCF_013301565.1 |
| Coprococcus eutactus MSK.18.32 | GCF_013299605.1 |
| Faecalicatena fissicatena MSK.9.3 | GCF_013300195.1 |
| **Lantibiotic resistance gene** | **HMM/Uniprot accession** |
| **Cell wall biosynthesis** | |
| murG | Prokka annotation |
| murJ | Prokka annotation |
| **Penicillin-binding proteins** | Prokka annotation |
| dltA | TIGR01734.1 |
| dltB | TIGR04091.1 |
| dltD | PF04914 |
| **Efflux pump** | |
| lanF | Prokka annotation |

| | |
|---|---|
| lanE | Prokka annotation |
| lanG | Prokka annotation |
| bceA | Prokka annotation |
| bceB | Prokka annotation |
| **Others** | |
| telA | PF05816 |
| mprF | Q2G2M2; Q8Y6I9 |
| nsr | P23648 |
| lanI | PF18218 |

**41 unique lanthipeptide candidates from the gut microbiome**

>WP_065541929.1_1408289 Blautia coccoides
MAKFDDFDLDVTKTAAGEGGVEPRITSKSLCTPGCVTGILMTCPVQTATCGCQITGK
>WP_117638663.1_224961 Dorea formicigenerans
MAKFDDFDLDVTKTAAQGGIEPKYKSKSVCTPGCPTGILMTCPLKTATCGCHITGK
>WP_119197961.1_129175 Blautia sp. AM47-4
MAKFDDFDLDVTKTAAQGGIEPKYKSKSACTPGCPTGILMTCPLKTATCGCHITGK
>WP_119197961.1_129369 Blautia sp. AM47-4
MAKFDDFDLDVTKTAARGGIEPKYKSKSACTPGCPTGILMTCPLKTATCGCHITGK
>WP_117994378.1_29961 [Ruminococcus] gnavus
MPNFDDFDLDLRENGDDSVMQTGVSITSDGFCDTDVITSLVSCTRNGQFATCMCSTCCTGHTAPAGCSTGC
>WP_090169706.1_68663 Pseudobutyrivibrio sp. 49
MEKFNNDFNLDVTVSKDDTKKADVKITSKSLCTPGCVTGVLMGCNNKTATCNCSVHVG
>WP_070088801.1_6782 Merdimonas faecis
LCGLKNSATSKQTVIEGEKIMGKMDDFDLDLRKIAENGNSANALSASDMITSEIISKVTETITRTFKGQCVSVETPTTGMTSAC
CKKGGTDVEPQCVP
>WP_090169706.1_68164 Pseudobutyrivibrio sp. 49
MEKFNNDFNLDVTVSKDDTKKADVKITSKSLCTPGCVTGLLMGCAGSSATCNCSVHVG
>WP_090169706.1_68413 Pseudobutyrivibrio sp. 49
MEKFNNDFNLDVTVSKDDAKKADVKITSKSLCTPGCVTGLLMGCAGSSATCNCSVHVG
>WP_101696280.1_129779 Clostridium minihomine
MYHRRKNIMGKFDDFDLDFTKVSAAAENSSERAGIAVPKLAINWSKISCACSPSDMTVCRAGAPGQLRC

>WP_124067899.1_52626 Clostridium sp. E02
MAKFDDFDLDLKATSTKSGGISPMVTSVSLCTPGCITGVIMTCTIKTATCGCHVAGK
>WP_125152452.1_480876 Clostridium sp. Marseille-P4200
LVTTYLKNILEGGKMGKLNDFDLDLKVKETTKKGVEPTWKSKSFCTPGCVTGILMTCTSNGCK
>WP_065541929.1_1408112 Blautia coccoides
MAKFDDFDLDVTRMESGNNKTQPRITSISLCTSGCKTAFFGTCSFTKDK
>WP_068555846.1_216193 Thermotalea metallivorans
MPKYDDFDLDIQVKKVDNLVEPNITSVSLCTPGTCWNTCGGTSTLNTNCCIATYLCSLAESCR
>WP_070088808.1_6842 Merdimonas faecis
MGKMDDFDLDLRKIAENGNSANALSASDMITSEIISKVTETITRTFKGQCVSVETPTTGMTSACCKKGGTDVEPQCVP
>WP_008429197.1_38472 Clostridium sp. LS
MREKNMGKLDDFDLDVKVKADTTKVGPAITSKSLCTPGCITGVLMCITQNSCVSCKSCIKC
>WP_068555846.1_215943 Thermotalea metallivorans
MPKYDDFDLDIQVKEVSNAAEPNITSVSLCTPGTCYQGCSGDTTFNSNCCPSNYTVCITKTCFTCA
>WP_118445440.1_623 [Ruminococcus] gnavus
MNKDFDLNLKADKSATEGPAPRITSVAYCTPGCVTGELCGSSECGLTRNCTGSILWA
>WP_090169706.1_68906 Pseudobutyrivibrio sp. 49
MAKLKNDFDLDLTVSKDNTKSADVQFKSISLCTPGCPTGILMGCHKCPSGSDTVYTK
>WP_085321196.1_7739 Clostridium botulinum
MGKFDDFDLDVNVNKGKEKQSRIYSCTPGTCARTCDEACAPNYTFGTNCCASFNICFFQ
>WP_118445440.1_405 [Ruminococcus] gnavus
MNKDFDLNLKADKSTTEGPAPRITSVAYCTPGCITGDLCGSSECCLTRNCTGSILCA
>WP_118445440.1_1056 [Ruminococcus] gnavus
MNKDFDLNLKTDKNTTEGSAPRITSVAYCTPGCITGELCGSSECGLTRNCTGSILCA
>WP_118445440.1_810 [Ruminococcus] gnavus
MKTVMKGEKKMNKDFDLNLKADKSATEGPAPRITRVAYCTPGCVTGELCGSSECGLTRNCTGSILCA
>WP_077834666.1_106377 Clostridium roseum
VSSLSIIVNVNMSYLYILKQIIIKGDIIMPKFDDFDLDVKINKGKGVKPQIATSAVACTPGSCWGCPETSTFASACCNVSDNCG
D
>WP_077852909.1_9492 Clostridium aurantibutyricum
VSSLSIIVNVNISYLYILKQIIIKGDIIMPKFDDFDLDVKINKGKGVKPQIATSAVACTPGSCWGCPETSTFASACCNVSDNCGD
>WP_118704912.1_5418 Ruminococcus sp. AM49-10BH

MANYDDFDLDIRKIKGNMEAEPKGTVAICLTTGTATTCTMPTLCDTSIVTLASCDGTCVGCTNTEKPCSANTCSACNSYCGG
ACRK
>WP_118662284.1_8946 Coprobacillus sp. AF13-4LB
MNTLKDFDVNLNSVNNDDGDDTSTYSLWTSATTVPCSVAASISALSAFSSNWTVTGDPNYTESKC
>WP_080549726.1_102266 Clostridium perfringens
MLKDFDLNLKMETGSSDDKEPRITSVAYCTPGCLTGELCGSSECGLTRSCTNTWLCS
>WP_016224249.1_1916777 Lachnospiraceae bacterium 3-2
MSNFKDFDLDLRNVASGGNAEPNGTTLVCVASKMTIKQKCNSVETPTTGMTSACCKKKNAADEPQCV
>WP_087213613.1_6459 Drancourtella sp. An177
MGNLNDFDLDLRKFAQNTEGEGRVDTGDVVSITTSITTIPCQLLSLYSVEKCGEPSNAAIPTTAMTVNCCTKEGGNVPNCV
>WP_119245824.1_43152 Ruminococcus sp. AM50-15BH
MAKFDDFDLDIIEKKVRRDIIPASITSQHSFCTPNCLTGFLCPPKTQLTCTCKLKGQ
>WP_117416421.1_13043 Clostridium sp. PI-S10-A1B
MNKDFDLDLKMDKGSVDVGESKIASIVYCTSGCLTGTTCETSECGITRSCTNTVLCY
>WP_118627011.1_3447 Clostridium sp. AM43-3BH
MGKYDDFDLDLKKVMSNNTASEAESVVSILQTLATECFTFDGSNTCNCPSTDCSPCDMTTSCRRADPQIMRC
>WP_118470785.1_29014 Coprobacillus sp. AF19-3
MKDVKLDDFDLDIVGGDDNENDDGVSPASSISTGWVSISVWISNGLSTNFTESKC
>WP_066580644.1_3898242 Clostridium sp. Marseille-P2538
LQYGGKAMGKYDDFDLNVQDENIGGDSSGAQPKGFWSEVGKSVVSSLVTSCTGNCPSVKCTGNCTSTCTTSMTTNCLCHR
R
>WP_117908709.1_956 Ruminococcus sp. AF32-2AC
MANYDDFDLDIRKIKGNMEAEPKGTAAICLTTGTATTCTMPTLCDTSIVTLASCDGTCVGCTNTEKPCSANTCSACNSYCGG
ACRK
>WP_119623224.1_62755 Lachnospiraceae bacterium GAM79
MGNYSDFDLDIRSGEPAEPAAITSKACEVLWEMATASIDYCGKISELICPTDACTVGCSSDTCSACHSYCGAACRRIG
>WP_106062361.1_54850 Clostridium liquoris
MPNFEDFDLDLKKVKTNSDKVSVNTPTPTISYLVHCNTADCPSTDCSPGDMTIGCYAADKGVCRC
>WP_105309223.1_429064 Dorea sp. Marseille-P4042
MKRKELIMGKYDEFDLDIKEVKANSSKGTARSTWGCVEKSIEISQFVTNALSCQSCGVCSAGGRSQCAPCTGSSKGSQARC
>WP_005342481.1_22836 Dorea formicigenerans
MKRKELIMGKYDEFDLDIKEVKANSSKGTARSTWGCVEKSIQISQFVTNALSCQSCGVCSAGGRSQCAPCTGSSKGSQARC
>WP_107030381.1_23895 Coprobacillus sp. AM17-34

MPNLDEFDLDPVVSTAENGNDGISPQSYSVTTKPCLVTDVISVLSQVSTSLSTNPGYTESKC

**Six unique class I lantibiotic sequences**

>BpSCSK BpcA1 Blauticin
MAKFDDFDLDVTKTAAGEGGVEPRITSKSLCTPGCVTGILMTCPVQTATCGCQITGK
>Dorea formicigenerans Lan-Df
MAKFDDFDLDVTKTAAQGGIEPKYKSKSVCTPGCPTGILMTCPLKTATCGCHITGK
>Pseudobutyrivibrio sp. 49_2 Lan-P49.2
MEKFNNDFNLDVTVSKDDTKKADVKITSKSLCTPGCVTGVLMGCNNKTATCNCSVHVG
>Pseudobutyrivibrio sp. 49_1 Lan-P49.1
MEKFNNDFNLDVTVSKDDTKKADVKITSKSLCTPGCVTGLLMGCAGSSATCNCSVHVG
>Clostridium sp. E02 Lan-CE02
MAKFDDFDLDLKATSTKSGGISPMVTSVSLCTPGCITGVIMTCTIKTATCGCHVAGK
>Blautia obeum NsoA1 Nisin O
MGKFDDFDLDVTKTAAQGGIEPKYKSKSACTPGCPTGILMTCPLKTATCGCHITGK
>NisA Nisin A
MSTKDFNLDLVSVSKKDSGASPRITSISLCTPGCKTGALMGCNMKTATCHCSIHVSK

**Codon optimized gene sequences for expression in *E. coli***

>BpcB, codon optimized
ATGAAGAAGCTGTTCTACGACATCGGCGAGTTCATGTATCGCCGTCCGACCGAGTACAAAAGCCAGATCGATTTCAGC
GAACACGAAGTCAAACTGATCTGCAGCAATCCGGCATTTCGCGAAAAAGTCAACATTGCGAGCCCGTCTCTGGTCGAA
ATGATGGACATCTACATGAAGAACCCGAAGCAGCTGAGCGAAAAGAAAAGCAACGGCCTGAACATCAGCCTGCTGAA
ATACCTGATCCGTAGCAAAGAACGTACCACCCCGTTTGGCTTATTTACCGGCGTTGGTACCGGTTGTTTCAGCAAAAGC
GAGAAGTTCCCGATCCTGATGACCAAGACCGAGAAGAAGGTCAACGTCGATTCTGAGTGGCTGTTTGGCCTGGTTAAC
ATCGTCGAGAAGAACTACGCGGAGAAACTGGAGTTCAAGTTCAACGACGCCTGCTACATCAAAGGCAATCGCGTCATC
CTGGTCTATAGCACCGAAAAAGACGCGGAAGAGATCAGCATCCGTTTCACCAAGGTCTTCAAGGTCCTGTTCGACAAG
ATCAAGGACTACGAGCGCTACGAGAAGTTCATCGAGATCCTGAGCAGCGAATACCCGTCCACCAGCATGAAAAAGATC
AAGCTGTACATCAACGAGCTGATCTCCAAGGGCTTCCTGATTAGCAACCTGCGTCCGAGTTTTAGCAACGCTGACCCGC
TGATGTACTTCATCAAGCAATGCGAGCGCATGGAGATTACCGATATCTGCGAGAAGGCGAACGAGATCTACAAGATGT
GCGAGGATTACAGCAAAACCGACATTGGCAACGGCATCATCAAGTACAACGCGATCAAGACCAAGATGCAGACCCTGT
ATAAGTGCAGCAGCTACCTGCAGGTAGATACCGTTATTGGCGGCGGCGATTTTCAACTGAACCAGGATATCAGCAAAG
CGATTTGCGAAGTTGCGTCTCTGTTCGTCTACCTGAGCAACAGCCCGAAAAAGCAGCACGGTTATCTGGAACATTACCG

CAACAAGTTCATCGAGAAGTACGGCATCGATCGCGAAGTTCCGCTGCTGGAAATGATCGATAGCAGTAACGGTATTGG
CGCACCGACCCCGTATTTAAAACCGCAGAACGACTTCTACGACGAGTACAACACCAAGGACAACTACAAGTACGAGCT
GAAGAACTACTTCCTGGTCGAGTACGAGAAAGCGCTGGCGAATAACAGCTACATCGACATCAACATGGAGACCCTGCA
GAAGATCACCGATTGCACCGTCCAGGAAGAAGAAATTCCGATCAGCCTGGAGCTGTACTTCATCCTGAAGGTCGAAGA
CGGCAAAGTTAGCCTGAATCTGAGTCCGAATTGCGGTAGCTTTGTTGCTGGTAAAACCTTCGGCCGTTTTAGCGTTCAG
AGCGATAACTTCGCGAACGTCCTGAAGAAGTGCAACAAAGAAGAACGCAAAATCCGTAGCCCGCACAGCGAAATCTG
CGAAATCAGCTTCCTGCCGAGTCCGACCCGTAACGGTAATATTGTTCGCACCCTGAGCTTTCGCGAAAAAGAAACCGCG
GTTTTTACCTGCGGCAGCAAAGATAAGAAGGACATCGTCAGCCTGAACGACATCTACATCGGCATCTTCAACGAGAAG
TTCTACGCGCGCGATAAGAAAACCGGCAAGCTGATCATCTTCGAGTCCAACAACATGTACAACCCGATGCTGAACCCG
AACGTCTTTCGCTTTCTGCAGGATATCAGCTACGAAGGTAAACGCGAGTGGTCCGAATTTCCGTGGAGCTACATCTACG
CAGATCTGCGCCATATTCCGACCATCAAGTACAAAGGCATCGTCCTGCAAAACGAGAAGTGGAAGGTCAACATCCAGG
AACTGGAACTGCTGAAGAAGGACTTCGAGTCGTTCAAAGAAAAATTCCTGGCGCTGATTATTGGCCGTAACATGCCGCT
GAACATCTACATTGTTGACGCGGATAACCGCATTCGTCTGAATCTGGCTACCGATCTGAGCATGCGTATTGTCTACGAC
GAGTTCAAGAAGCACAAGGACAGCGATCTGGTCTTCGAGAAAGTCGAAAACGGCAGCGACATCATCTACGATGACGGC
AAAACCTACGCAACCGAAATCGTTGTCCCGCTGTTCCGCAAAAACAAAGAAGAACTGAGCCTGATTCCGCTGAGTCAA
AAGACCTACACCCGCGAACAGCACATGATTCTGCCGTTCAACAACTGGCTGTACCTGAAGCTGTACTGCAACGAAAAC
CGCGAAGAAGAGCTGATCGCCTTCTACATCATGGACTTCTACGAGAGCCTGAAAGAGAAATACGGCATCTCCTACTTCT
ACATGCGCTACGCGGATCCGAAACCGCATATTCGTCTGCGTTTACACGCAACCCGCGAATTACTGCTGCAGGTTTATCC
GCAAATCCTGAAGTGGTACAGCGAACTGTTCAGCGATCAGATTGTCGGCGATATGACCATCAGCGTCTACGATCGCGA
AATTGAACGTTACGGCGGCGCTTTACTGATGGATACCGCGGAAAAAGTCTTCTTCGAGGACAGCTACATCGTCGAAAA
CATCCTGCGTCTGAAACGTCTGGGCAAAATTAGCCTGAACCTGGACGACGTTGCAGTTGTCTCCATCATCATGTACGTC
AGCCAGTTCTACAACAAGTACGAAGAACAGCTGCAGTTCCTGAGCATTAACTACCACAGCAGCGACTTCATCAGCGAG
TTCAAGAAGAAGAAGGACAACCTGCTGCGCATCTGCGATATCGAAAACGAGTGGAACAACCTGAACAGCATGAAGGA
CGGCAAGGTTGTCTACGAACTGATGTGGCGTCGTTGCAAAGTCATCAGCGAGTACAGCGAGAAGATCCGCAAAATCAA
CCCGGATCCGATGTTCAAGAACAGCATTGTCGCGAGCGTCATCCATCTGCATTGCAACCGTCTGATCGGTACCAATCGC
GAACTGGAACGCAAACTGATGGCATTCGCGGAAAGCGTTCTGTACGCGAAAAAGTACGTCATGCGCCGTATCGAAGTT
AACGGCAAAAAGTAA

>BpcC, codon optimized

ATGGACAACAGCATCATCAGCATCGTCAAAGAGATTGCGCGTAACCTGGGCGATTACGATAACGTCAAACGCATCGTC
GCGGACAAAGATAACATCCGCGTCATGGGCGAGTTCAACTTTCAAGGTTGGGAACCGCTGACCCTGAGTCACGGTATT
CCGGGTATTTGTCTGCTGTACGGCAAACTGATGGAGTGCTTTCCGGACGAAGAAATCTGGGCAGAACTGGCACATCAGT
ATCTGGGGTATCTGGTCCAGGAGATCAACAAAGAGGGCTTTCAAACCCTGAGCATGTTTAGCGGTACCTCTGGTATTGG
TCTGGCAGTTGCATCAGTCAGCAACAACTTCTGCAACTACAACAAACTGCTGAACACCATCAACAGCTACATCATCAAC

TGGTTCGACGAGTTCATCGACAGCATCGATCTGAAAAAAGGCACCCGTAGCATCTGCTACGACGTTATTGAAGGCCTGA
GCGGCATTCTGAGCTATTGCAGCATCTACTACGAGCAGGAAAGCTTTAGCCCGATTCTGCTGAAAGGCCTGAAAAAACT
GGTCGAGCTGACCTACGACATCGAAGTCAAAGGCTATCACGTTCCGGGTTGGTATATTCCGAGCGACAACCAGTTCAGC
ACCGTTGAAAAAGAGCTGTACCCGTACGGTAACTTCAACACCAGCTTCAGCCACGGCATTGCAGGTCCGCTGACCCTGC
TGAGCGAAATGAAATCCAAAGGCTTCATGATCGAGGGCCAGGAAGAAGCGATCGAGAAAATCGTCAAATTCCTGTTCG
ACTTCCGCAGCAACGATCAGAAACGCGACTTTTGGAAAGGCCAGATCGATTTCCACGAATACATCACCGGCAAAGTCA
GCGAGAAAAACATCATCCGTCGCGACGCTTGGTGTTACGGTAATCCGGGCGTTTGCTATTCTCTGATCATGGCTGGCAA
CGCGATGAAAAACCAGAGCTGGATCGACTACGGCATCCACAACATGAAAAAGACCCTGAGCGACGTCAAAGGCATCTT
TAGCCCGACCTTTTGCCACGGTTTTTCCGGTCTGTACCAGGTTGTCAACAGCATCGAATTCACCATCGGCAAAGACATCT
TCTACAGCGAGAAAAAAGAGCTGCTGAACAAAATCATGAGCTTCTACGACAGCAACTACATCTTCGGCTTCCGCAACA
TGGAAGTAGGCGACGAAAACGGCAACATTCGCGCATTTGAACATCTGGGTCTGCTGGACGGTACCATTGGCGTTTGTCT
GGCACTGCTGGAAGGCGAACACAAAACCAAAAACATCTGGAAACGCGCATTTCTGCTGGCATAA
